# Parallel recruitment pathways contribute to synaptonemal complex assembly during mammalian meiosis

**DOI:** 10.1101/2022.04.14.488335

**Authors:** James H. Crichton, James M. Dunce, Willy M. Baarends, Owen R. Davies, Ian R. Adams

**Author notes:** **Authors for Correspondence** Ian Adams, Owen Davies. These authors contributed equally to this work (James H. Crichton, James M. Dunce).

## Abstract

During meiosis, the synaptonemal complex (SC) assembles between paired chromosomes, binding them together in close apposition, and facilitating recombination. SC assembly is thought to occur through the hierarchical zipper-like recruitment of axial elements, followed by transverse filaments and then central elements. However, the rapidity of SC formation in mammals has hitherto hindered investigation of its assembly mechanisms and their relationship with recombination. Using super-resolution imaging of separation-of-function mouse mutants, we show that, contrary to the hierarchical assembly model, central element protein SYCE2 is recruited to recombination sites early in SC assembly, and independently of SYCP1-containing transverse filaments. Further, SYCE2-TEX12 binds DNA *in vitro*, and SYCE2-containing bridges physically link paired chromosomes at recombination sites prior to transverse filament recruitment and chromosome synapsis. These data suggest that mammals integrate parallel recruitment pathways to assemble a mature SC: one recruiting central element proteins to recombination sites, and another recruiting transverse filaments to chromosomes.

## Introduction

Meiosis reduces the number of chromosomes during gametogenesis to allow diploid germ cells to mature into haploid gametes. In mammals, reductional meiotic division is achieved by recombination-mediated pairing of homologous chromosomes, followed by maturation of a subset of recombination sites into crossovers, and then segregation of homologous chromosomes between daughter cells. The synaptonemal complex (SC) is a supramolecular protein structure that synapses together paired homologous chromosomes, and is required for the progression of recombination and crossover formation (Hunter, 2015; Kouznetsova et al., 2011; Zickler and Kleckner, 2015). SC assembly is initiated at crossover-fated recombination sites in *Sordaria* and budding yeast, but independently of recombination at centromeres/pairing centres in budding yeast, *Drosophila* and *C. elegans* (Cahoon and Hawley, 2016; Gao and Colaiácovo, 2018; Zickler and Kleckner, 2015). SC assembly therefore likely involves distinct mechanisms in different species (Cahoon and Hawley, 2016), and SC assembly mechanisms in mammals and their relationship with recombination remain poorly understood.

In mammals, the SC is thought to assemble hierarchically (Figure 1A) (Cahoon and Hawley, 2016; Fraune et al., 2012; Gao and Colaiácovo, 2018). First, axial/lateral elements containing SYCP3 assemble along chromosome axes. Chromosome pairing, mediated by meiotic recombination from sites of programmed DNA double strand breaks (DSBs), then generates regions of paired chromosome axes that are separated by ∼400 nm. These are brought into ∼100 nm synapsis through recruitment of transverse filament protein SYCP1, arranged with the N-termini of its coiled-coil structure within the SC’s central element (CE), lying midway between synapsed axes, and its C-termini within chromosome-bound lateral elements (Figure 1A) (de Vries et al., 2005; Dunce et al., 2018; Schücker et al., 2015; Zickler and Kleckner, 2015)). SYCP1 initially assembles into a tetramer lattice that can support only short-range contacts between chromosomes (Figure 1A) (Crichton et al., 2022; Dunce et al., 2018). Then, recruitment of CE protein SYCE3 disrupts the SYCP1 tetramer lattice, and remodels it into an integrated SYCP1-SYCE3 lattice that relies on SYCE3 self-assembly interactions for its structure (Figure 1A) (Crichton et al., 2022). SYCE3 recruits CE complexes SYCE1-SIX6OS1 and SYCE2-TEX12, which confer stability to the integrated SYCP1-SYCE3 lattice, and enable its long-range extension along the entire chromosome axis to achieve full synapsis (Figure 1A) (Crichton et al., 2022). In mice, SC assembly occurs at different regions of the chromosomes asynchronously, such that in zygotene nuclei, some regions are still assembling axial/lateral elements whilst others are fully synapsed with a mature SC.

**Figure 1.**
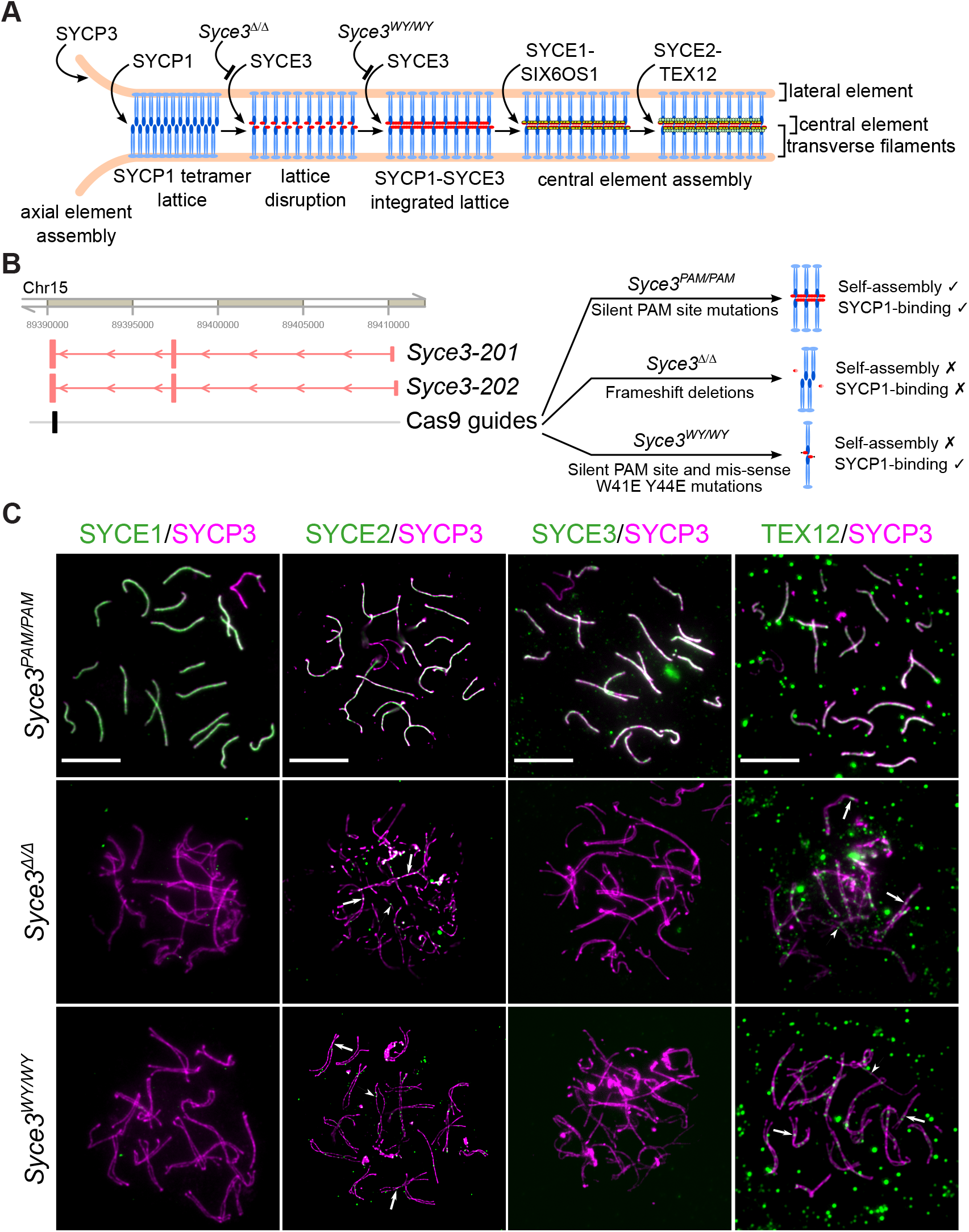
Central element proteins SYCE2 and TEX12 are recruited to meiotic chromosomes in *Syce3* mutant spermatocytes. (**A**) Schematic diagram outlining the hierarchical model of mammalian synaptonemal complex (SC) assembly. SYCP3 (orange) is recruited to the axial elements of the assembling SC. The transverse filament component SYCP1 (blue) then assembles between two axial elements that are in close proximity to form a tetrameric lattice linking two synapsing chromosomes. SYCP1 recruits SYCE3 (red), which disrupts the SYCP1 lattice. SYCE3 self-assembly interactions then allow an integrated SYCP1-SYCE3 lattice to form. SYCE3 is required for the subsequent recruitment of the interacting central element (CE) proteins SYCE1 and SIX6OS1 (yellow), which are in turn required for recruitment of the interacting CE proteins SYCE2 and TEX12 (green). (**B**) Outline of *Syce3* mouse mutants used in this study. The structure of the mouse *Syce3* locus (mm10 genome co-ordinates), its predicted transcripts (Gencode VM18), and location of Cas9 guides used to mutate this locus are indicated on the left. The two *Syce3* transcripts encoded at this locus differ in their 5’ non-coding exons, but encode identical proteins. The *Syce3* mutations used in this study and their predicted effects on SC assembly and SYCE3 protein interactions are indicated on the right. *Syce3*^*WY/WY*^ mice carry mis-sense mutations in the SYCE3 self-assembly interface (W41E, Y44E) in addition to silent mutations in the PAM site of the Cas9 guides used to introduce these mutations. *Syce3*^*PAM/PAM*^ control mice contain the same silent mutations in the PAM site that were used to construct the *Syce3*^*WY/WY*^mice, but not the mis-sense mutations. *Syce3*^*Δ/Δ*^ mice possess frameshift deletions upstream of the W41, Y44 residues mutated in *Syce3*^*WY/WY*^ mice. (**C**) Widefield imaging of meiotic chromosome spreads from control pachytene *Syce3*^*PAM/PAM*^ and asynapsed pachytene *Syce3*^*Δ/Δ*^ and *Syce3*^*WY/WY*^ spermatocytes were immunostained for SYCP3 (magenta) and the indicated central element components (green). Example weak SYCE2 and TEX12 microfoci associated with the potentially synapsed or asynapsed regions of SYCP3 axis are indicated with arrows and arrowheads respectively. Scale bar 10 µm.

This hierarchical model for mammalian SC assembly is consistent with phenotypic analyses of mice harbouring mutations in SC components. SC assembly progresses to the stage at which the mutant protein would assemble, but downstream SC proteins are not recruited, resulting in defective SC assembly, chromosome asynapsis, spermatocyte death and male infertility (Bolcun-Filas et al., 2009, 2007; de Vries et al., 2005; Gómez-H et al., 2016; Hamer et al., 2008, 2006; Schramm et al., 2011). It is less clear how SC assembly might incorporate interactions between the SC and recombination sites, either to nucleate assembly, or to stabilise and/or regulate meiotic recombination. Some short stretches of SC that assemble in *Syce1* mutant mouse spermatocytes overlap with recombination foci (Bolcun-Filas et al., 2009), but there does not appear to be an obligate association between SC assembly sites and recombination sites in wild-type mouse spermatocytes (Gruhn et al., 2016). Further, it is uncertain whether interactions between SC components and recombination proteins reflect roles in recruiting SC to recombination sites during its assembly, roles regulating recombination in the fully assembled SC, or both (Bolcun-Filas et al., 2009; Tarsounas et al., 1999; Yang et al., 2008).

We previously generated mice carrying mutations that target the two stages of SYCE3-mediated remodelling of the SYCP1 tetramer lattice (Figure 1A,B) (Crichton et al., 2022). Firstly, the *Syce3* WY separation-of-function mutation (*Syce3*^*WY/WY*^) binds to SYCP1 and disrupts SYCP1 tetramer lattices, but cannot self-assemble, so fails to form integrated SYCP1-SYCE3 lattices (Figure 1A) (Crichton et al., 2022). Secondly, *Syce3* truncating frameshift deletions (*Syce3*^*Δ/Δ*^) resembles a *Syce3* null mutation (Schramm et al., 2011), so cannot remodel SYCP1 tetramer lattices, which remain intact (Figure 1A) (Crichton et al., 2022). SYCP1 is recruited to meiotic chromosomes, and forms foci on asynapsed and synapsed regions of the chromosomes in both *Syce3*^*WY/WY*^ and *Syce3*^*Δ/Δ*^ spermatocytes. However, in accordance with their targeted actions, SYCP1 foci are far fewer in number, and immunostain much more weakly, in *Syce3*^*WY/WY*^ than *Syce3*^*Δ/Δ*^ spermatocytes (Crichton et al., 2022). Hence, these *Syce3* mutants trap different stages of SC formation, providing a unique opportunity to probe the molecular basis of SC assembly and its relationship with recombination.

Here, we use super-resolution imaging of *Syce3* mutants to show that, contrary to the hierarchical SC assembly model, CE proteins are recruited to meiotic chromosomes independently of SYCP1, localise to recombination sites prior to synapsis, and assemble into bridge-like structures that link paired pre-synaptic chromosomes at recombination sites. Our data suggest that mammalian SC assembly is achieved through the integration of multiple recruitment pathways operating in parallel, and that direct interactions between CE proteins and recombination intermediates can direct a subset of SC assembly events to recombination sites.

## Results

### SYCE2 and TEX12 are recruited to meiotic chromosomes in *Syce3*^*Δ/Δ*^and *Syce3*^*WY/WY*^ spermatocytes

It was previously reported that CE element proteins fail to be recruited to meiotic chromosomes in *Syce3*^*-/-*^ spermatocytes (Schramm et al., 2011). We therefore analysed CE protein recruitment in *Syce3*^*Δ/Δ*^ and *Syce3*^*WY/WY*^ asynapsed pachytene spermatocytes by widefield imaging. As expected, CE proteins SYCE1, SYCE2, SYCE3 and TEX12 were all detected on synapsed chromosomes in pachytene nuclei from control mice generated using CRISPR reagents that introduce silent mutations into *Syce3* (*Syce3*^*PAM/PAM*^, Figure 1B, Figure 1C) (Crichton et al., 2022). We did not detect SYCE3 on meiotic chromosomes in either *Syce3*^*Δ/Δ*^ or *Syce3*^*WY/WY*^ spermatocytes (Figure 1C). The lack of detectable SYCE3 on *Syce3*^*WY/WY*^ spermatocyte chromosomes is consistent with our previous findings that SYCE3 WY disassembles SYCP1 tetramer lattices into individual 2:1 SYCP1:SYCE3 complexes, which likely lack the cooperativity necessary to associate with chromosomes (Crichton et al., 2022). In agreement with the reported *Syce3*^*-/-*^ phenotype (Schramm et al., 2011), we were unable to detect SYCE1 on meiotic chromosomes in either *Syce3*^*Δ/Δ*^ or *Syce3*^*WY/WY*^ asynapsed pachytene nuclei (Figure 1C). Surprisingly, we did detect low-intensity axis-associated SYCE2 microfoci in both *Syce3*^*Δ/Δ*^ and *Syce3*^*WY/WY*^ mutant asynapsed pachytene nuclei (Figure 1C). These SYCE2 microfoci were present in both asynapsed and potentially synapsed regions of axes (Figure 1C). In accordance with their known existence in a co-folded SYCE2-TEX12 complex (Davies et al., 2012; Dunce et al., 2021), and despite the higher background and diffuse off-axis staining of anti-TEX12 antibodies, we also detected TEX12 microfoci associated with asynapsed and potentially synapsed regions of axes in both *Syce3*^*Δ/Δ*^ and *Syce3*^*WY/WY*^ mutants (Figure 1C). Thus, somewhat unexpectedly, weak foci containing central element components SYCE2 and TEX12, but not SYCE3 or SYCE1, are recruited to meiotic chromosomes in both *Syce3*^*Δ/Δ*^ and *Syce3*^*WY/WY*^ spermatocytes.

To test whether SYCE2 microfoci detected by widefield imaging might represent *bona fide* recruitment to meiotic chromosomes, we quantitatively analysed SIM super-resolution imaging of SYCE2 immunostaining in asynapsed pachytene *Syce3*^*Δ/Δ*^ and *Syce3*^*WY/WY*^ spermatocytes (Figure 2A). Computational simulations strongly suggest that the association of weak SYCE2 microfoci with meiotic chromosome axes does not represent chance overlap: SYCE2 microfoci were significantly located closer to SYCP3 axes than might be expected from randomly distributing these foci within the nucleus (Figure 2B). Furthermore, we used the distribution of the distances between SYCE2 microfoci centroids and the SYCP3 axis mask to assign a 140 nm threshold distance for axis association of SYCE2 microfoci (Figure 2B). Approximately seventy axis-associated SYCE2 microfoci were detected per asynapsed pachytene nucleus in these mutants, significantly more than would be expected from chance axis-associations obtained from randomly distributing these foci within the nucleus (Figure 2C). Thus, SYCE2 microfoci are recruited to meiotic chromosome axes in *Syce3*^*Δ/Δ*^ and *Syce3*^*WY/WY*^ mutant spermatocytes. These findings differ from previous analyses of *Syce3*^*-/-*^ spermatocytes (Schramm et al., 2011), which could reflect differences in the *Syce3* alleles used, or advances in the sensitivity of imaging technologies.

**Figure 2.**
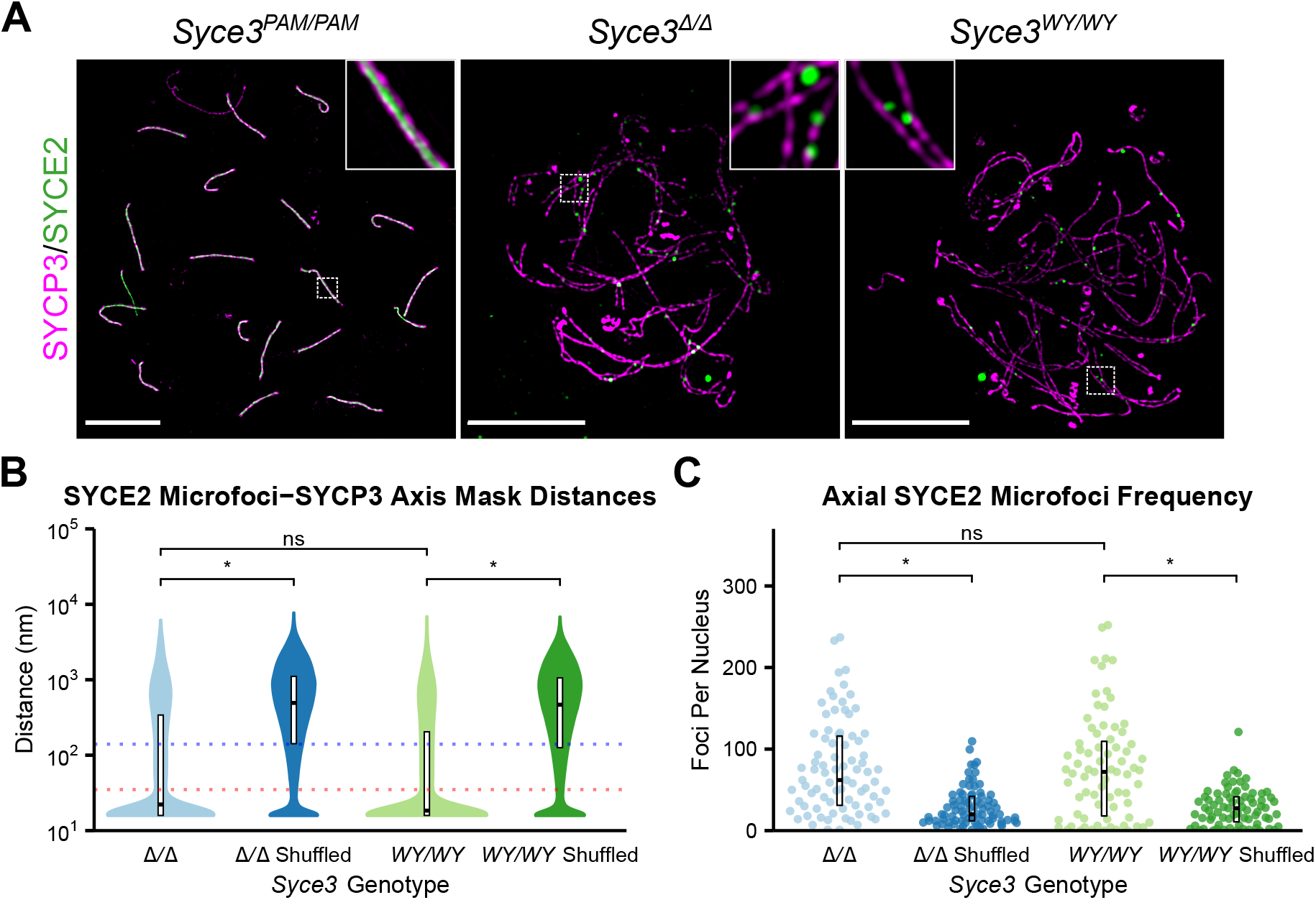
SYCE2 microfoci are enriched on chromosome axes in *Syce3* mutant spermatocytes. (**A**) SIM super-resolution imaging of *Syce3* mutant meiotic chromosome spreads immunostained for SYCP3 (magenta) and SYCE2 (green). Scale bar, 10 µm. (**B**) Distances from the centroids of each SYCE2 microfocus to the nearest point on the SYCP3 axis mask in *Syce3* asynapsed pachytene nuclei. Shuffled datasets were generated by assigning each SYCE2 microfocus centroid a random location within that nucleus for twenty repetitions. Blue and red dotted lines represent 140 and 35 nm; crossbars represent quartiles; *, p<0.01 (Mann-Whitney U test, paired test used to compare observed with shuffled datasets, nuclei medians are 18.4,445.6, 23.3 and 476.7 nm, n=81, 75 nuclei); 3 animals analysed for each *Syce3* genotype. Summary statistics are in Supplementary Table ST2. (**C**) Quantification of axis-associated SYCE2 microfoci in *Syce3* asynapsed pachytene nuclei. SYCE2 microfoci centroids within 140 nm of the SYCP3 axis mask were classed as axis-associated. Axial microfoci in shuffled datasets represent the median after twenty rounds of shuffling. Crossbars represent quartiles; *, p< 0.01; ns, no significant difference (Mann-Whitney U test, paired test used to compare observed with shuffled datasets, medians are 62, 20, 72 and 27.5 foci, n=81, 75 nuclei); 3 animals analysed for each *Syce3* genotype. Summary statistics are in Supplementary Table ST2.

### SYCE2 is recruited to meiotic chromosomes independently of SYCP1

As SYCP1 is recruited to meiotic chromosomes in *Syce3*^*Δ/Δ*^ and *Syce3*^*WY/WY*^ mutant spermatocytes (Crichton et al., 2022), we reasoned that these axis-associated SYCE2 microfoci could reflect recruitment of SYCE2 to SYCP1-containing structures, and formation of short CE-containing SC-like assemblies. However, SIM super-resolution imaging showed that most axial SYCE2 microfoci did not co-localise with SYCP1 foci (Figure 3A, Figure 3B). Approximately half of the axial SYCE2 microfoci in *Syce3*^*Δ/Δ*^ spermatocytes, and quarter of the axial SYCE2 microfoci in *Syce3*^*WY/WY*^ spermatocytes, overlapped with SYCP1 foci representing modest enrichments over the proportions expected by chance (Figure 3B asterisks, Figure 3C). As the extent of overlap between foci can be affected by staining intensity, we confirmed these findings by measuring distances between the centroids of axis-associated SYCE2 microfoci and their nearest axis-associated SYCP1 foci (Supplementary Figure S1A). Consistent with the overlap analysis, axial SYCE2 microfoci centroids are slightly closer to SYCP1 foci centroids than expected by chance, but are typically located too far away to co-localise (∼300 nm and ∼600 nm apart in *Syce3*^*Δ/Δ*^ and *Syce3*^*WY/WY*^ spermatocytes, respectively; Supplementary Figure S1A). Thus, although there is some association between SYCE2 microfoci and SYCP1 foci, the distribution of SYCE2 microfoci on chromosome axes is largely independent of SYCP1 foci.

**Figure 3.**
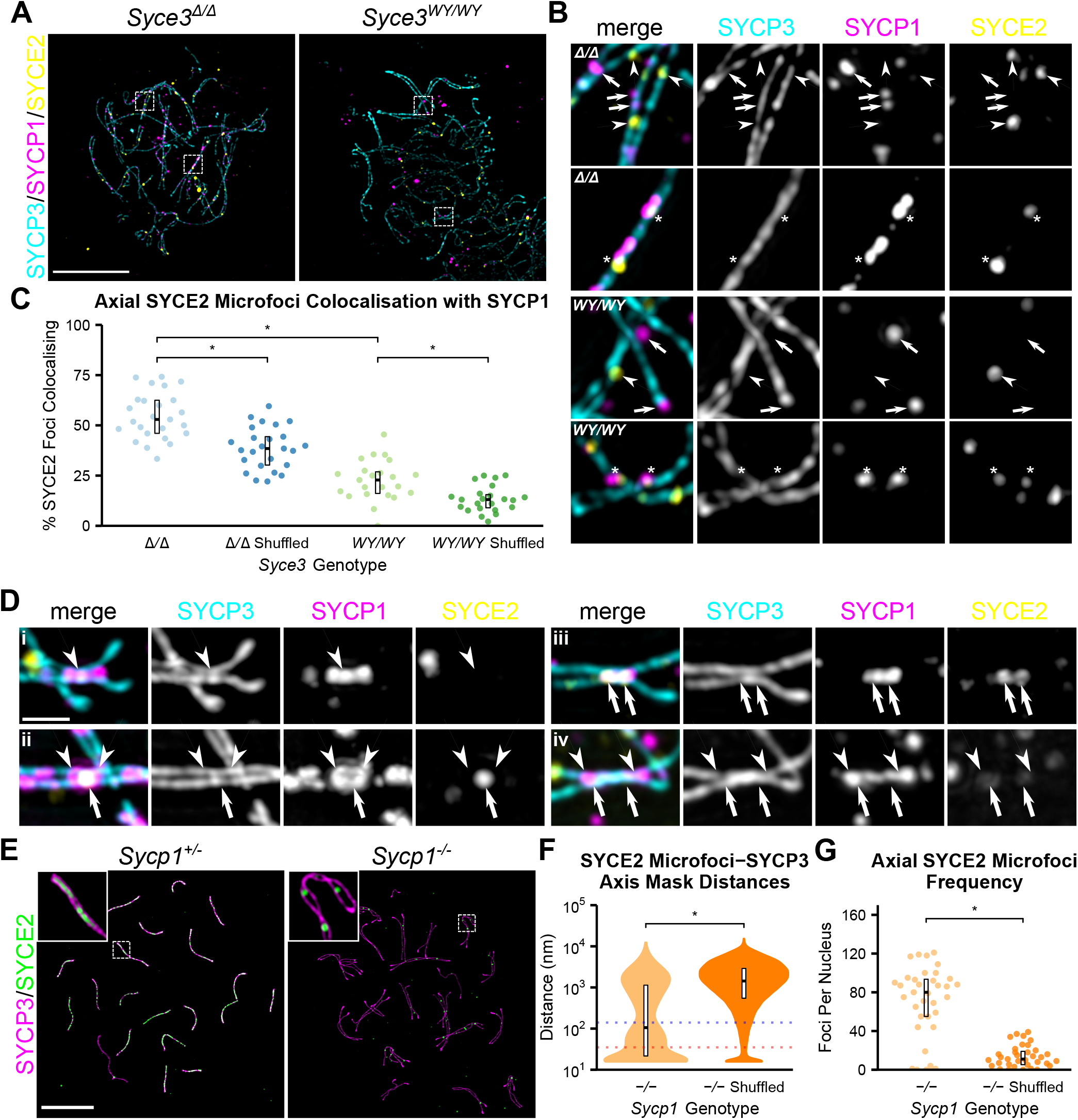
SYCE2 can be recruited to meiotic chromosomes independently of SYCP1. (**A**) SIM super-resolution images of asynapsed pachytene *Syce3* mutant spermatocyte chromosome spreads immunostained for SYCP3 (cyan), SYCP1 (purple) and SYCE2 (yellow). Boxed regions are shown at higher magnification in (**B**). Scale bar 10 µm. (**B**) Higher magnification images represent boxed regions in (**A**). Arrows indicate SYCP1 foci, arrowheads indicate SYCE2 microfoci and asterisks indicate co-localising SYCP1 and SYCE2 microfoci. *Syce3* genotype is indicated in the top right of the merged panel. (**C**) Colocalisation of axial SYCE2 microfoci and SYCP1 foci. The percentage of axial SYCE2 microfoci that overlapped an axial SYCP1 focus was calculated for each nucleus, and compared with the median percentage from shuffled datasets assigning axial SYCP1 foci to random positions on the SYCP3 axis for twenty repetitions. SYCE2 and SYCP1 foci overlapping by at least 1 pixel were classed as colocalising. Crossbars represent quartiles; *, p< 0.01 (Mann-Whitney U test, paired test used to compare observed with shuffled datasets, medians are 52.9, 38.5, 22.8 and 13%, n=25, 23 nuclei); 3 animals analysed for each *Syce3* genotype. Summary statistics are in Supplementary Table ST2. (**D**) SIM super-resolution images of large extended SYCP1 foci (magenta, arrowheads) associated with potentially synapsing SYCP3 axes (cyan) in asynapsed pachytene *Syce3*^*Δ/Δ*^ spermatocytes not co-localised (i) or co-localised (ii-iv) with SYCE2 microfoci (yellow, arrows). Scale bar 1 µm. (**E**) SIM super-resolution images of meiotic chromosome spreads from pachytene *Sycp1*^*+/-*^ and asynapsed pachytene *Sycp1*^*-/-*^ spermatocytes immunostained for SYCP3 (magenta) and SYCE2 (green). Boxed regions are magnified in insets. Scale bar, 10 µm. (**F**) SYCE2 microfoci-SYCP3 axis mask distances in asynapsed pachytene *Sycp1*^*-/-*^ spermatocytes. Distances between the centroids of SYCE2 microfoci and the SYCP3 axis mask and shuffled datasets were determined as described in Figure 2B. *, p< 0.01 (paired Mann-Whitney U test; medians are 89 and 1852 nm, n=35 nuclei); 2 *Sycp1*^*-/-*^ animals analysed. Summary statistics are in Supplementary Table ST2. (**G**) Axial SYCE2 microfoci frequency in asynapsed pachytene *Sycp1*^*-/-*^ spermatocytes. Axis-associated SYCE2 microfoci and shuffled datasets were defined as described for Figure 2C. *, p< 0.01 (paired Mann-Whitney U test; medians are 80 and 11 foci, n=35 nuclei); 2 *Sycp1*^*-/-*^ animals analysed. Summary statistics are in Supplementary Table ST2.

We next investigated whether the SYCE2 microfoci that did co-localise with SYCP1 in *Syce3*^*Δ/Δ*^ spermatocytes might represent assemblies of short CE-like structures. SYCP1 assembles into larger structures in *Syce3*^*Δ/Δ*^ than *Syce3*^*WY/WY*^ spermatocytes (Crichton et al., 2022) (Supplementary Figure S1B), but SYCE2 was not present in filamentous CE-like assemblies even in regions of *Syce3*^*Δ/Δ*^ spermatocytes that contained large extended SYCP1 foci linking paired proximate SYCP3 axes (Figure 3D). Hence, these co-localising SYCE2 and SYCP1 foci do not represent short stretches of CE-containing SC-like assemblies. Instead, they may represent chance association, trapped SC assembly intermediates, or dynamic attempts at assembling an SC that is unstable in the absence of SYCE3. Nevertheless, any interactions between SYCE2 and SYCP1 in these co-localised regions do not require functional SYCE3.

Whilst most SYCE2 microfoci did not co-localise with SYCP1, it remained possible that they were the remnants of a degenerating or unstable structure that was originally recruited by SYCP1. Hence, we directly tested whether SYCP1 is required to recruit SYCE2 microfoci to meiotic chromosomes by analysing *Sycp1*^*-/-*^ spermatocytes. In asynapsed pachytene *Sycp1*^*-/-*^ nuclei, we detected axis-associated SYCE2 microfoci that were substantially enriched relative to chance association (Figure 3E, Figure 3F, Figure 3G). Thus, SYCE2 microfoci can be recruited to meiotic chromosome axes independently of SYCP1. Given that SYCP1 is recruited to meiotic chromosomes in *Syce2* mutant spermatocytes (Bolcun-Filas et al., 2007), these data suggest that, contrary to current models of hierarchical SC assembly (Fraune et al., 2012), parallel assembly pathways are operating to recruit SC proteins SYCP1 and SYCE2 to chromosomes during mammalian meiosis.

### SYCE2 microfoci associate with RAD51 recombination sites

Given the association between SC assembly and recombination in budding yeast and *Sordaria* (Cahoon and Hawley, 2016; Gao and Colaiácovo, 2018; Zickler and Kleckner, 2015), we wondered whether SYCP1-independent recruitment of SYCE2 might represent its association with recombination sites. Accordingly, SIM super-resolution imaging of *Syce3*^*Δ/Δ*^ and *Syce3*^*WY/WY*^ mutants revealed that SYCE2 microfoci frequently co-localise with RAD51 recombination foci (Figure 4A). Further, the distances between axial SYCE2 microfoci and their nearest RAD51 foci were closer than expected from random shuffling along SYCP3 axes (Figure 4B; populations at approximately 120 nm and 80 nm were enriched in *Syce3*^*Δ/Δ*^ and *Syce3*^*WY/WY*^ mutants, respectively). The difference in SYCE2-RAD51 distances between *Syce3* genotypes may relate to direct or indirect effects of these mutations on the frequency, dynamics and/or progression of meiotic recombination (Supplementary Figure S2A). These findings in *Syce3* mutant spermatocytes are consistent with the co-localisation observed between SYCE2 and flares of RAD51 emanating from synapsed chromosome axes (Supplementary Figure S2B).

**Figure 4.**
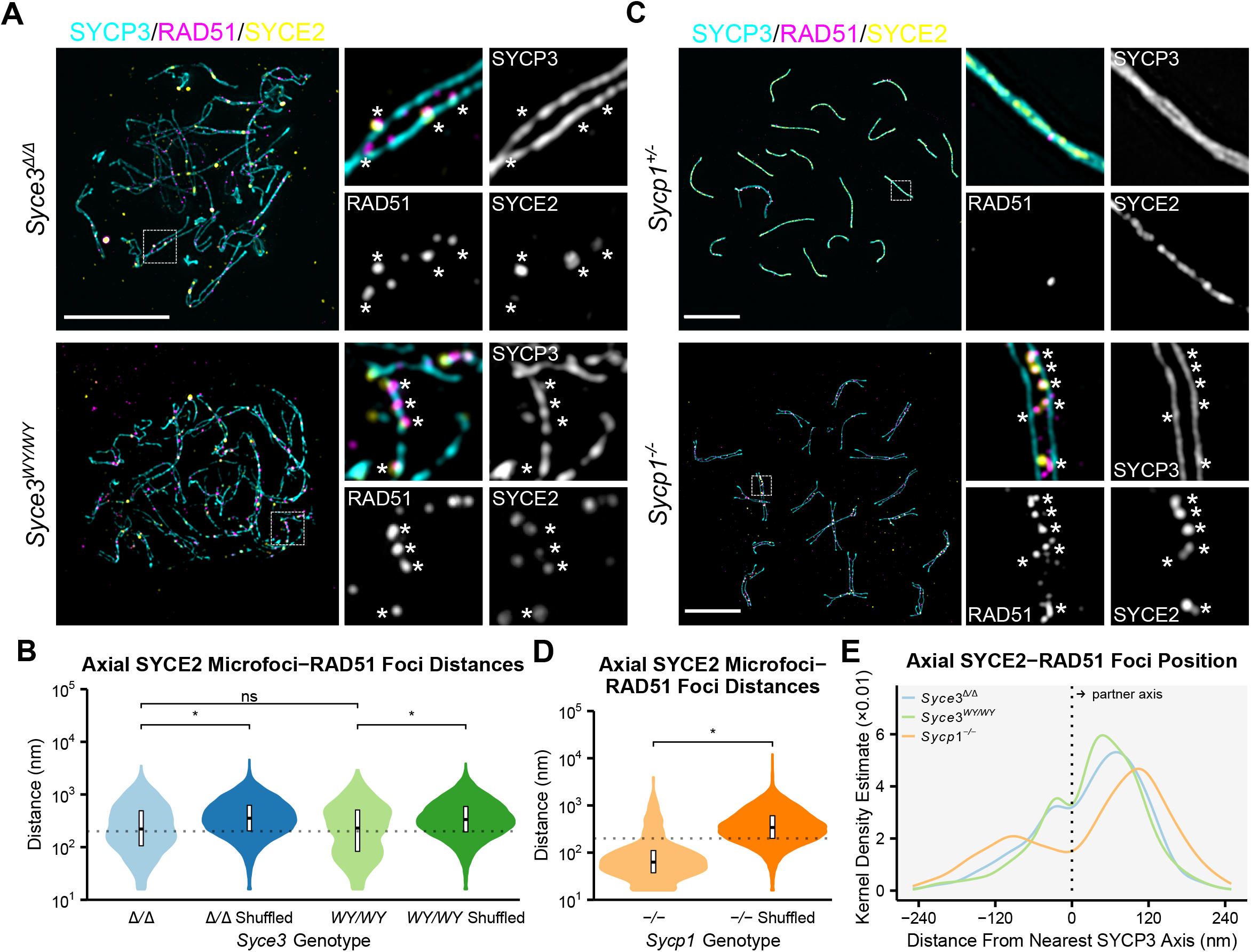
SYCE2 microfoci co-localise with RAD51 recombination sites. (**A**) SIM super-resolution images of asynapsed pachytene *Syce3*^*Δ/Δ*^ and *Syce3*^*WY/WY*^ meiotic chromosome spreads immunostained for SYCP3 (blue), RAD51 (magenta) and SYCE2 (yellow). Scale bar, 10 µm. (**B**) SYCE2-RAD51 foci distances in asynapsed pachytene *Syce3*^*Δ/Δ*^ and *Syce3*^*WY/WY*^ spermatocytes. Distances between the centroid of each SYCE2 microfocus that overlaps the SYCP3 axis and its nearest RAD51 focus centroid are shown alongside shuffled datasets assigning SYCE2 centroids to random positions on the SYCP3 axis for twenty repetitions. The gray dotted line marks 200 nm; crossbars represent quartiles; *, p<0.01; ns, not significant (Mann-Whitney U test, paired test used to compare observed with shuffled datasets, medians are 206.1, 367.7, 191.3, and 336.9 nm, n=56, 46 nuclei); 3 animals analysed for each *Syce3* genotype. *Syce3*^*Δ/Δ*^ and *Syce3*^*WY/WY*^ datasets are bimodal with additional populations at ∼ 150 nm and ∼80 nm respectively. Summary statistics are in Supplementary Table ST2. (**C**) SIM super-resolution images of synapsed pachytene *Sycp1*^*+/-*^ and asynapsed pachytene *Sycp1*^*-/-*^ meiotic chromosome spreads immunostained for SYCP3 (blue), RAD51 (magenta) and SYCE2 (yellow). Scale bar, 10 µm. (**D**) SYCE2-RAD51 foci distances in synapsed pachytene *Sycp1*^*+/-*^ and asynapsed pachytene *Sycp1*^*-/-*^ spermatocytes. Distances were determined as for (**B**). The gray dotted line marks 200 nm; crossbars represent quartiles; *, p<0.01; ns, not significant (paired Mann-Whitney U test, medians are 60 and 349.7 nm, n=31 nuclei); 2 animals analysed for each *Sycp1* genotype. The *Sycp1*^*-/-*^ dataset is bimodal with an additional population at ∼80 nm. Summary statistics are in Supplementary Table ST2. (**E**) Kernel density estimate that the SYCE2 centroids in axis-associated SYCE2-RAD51 foci are located at the indicated distance from their nearest SYCP3 axis trace in *Syce3*^*Δ/Δ*^, *Syce3*^*WY/WY*^ and *Sycp1*^*-/-*^ asynapsed pachytene spermatocytes. SYCE2 centroids within 250 nm of the SYCP3 axis trace and 200 nm of a RAD51 focus were classed as co-localised axial SYCE2-RAD51. Paired SYCP3 axes separated by less than 100 nm were excluded. 3 *Syce3*^*Δ/Δ*^, 3 *Syce3*^*WY/WY*^ and 2 *Sycp1*^*-/-*^ animals analysed.

We next tested whether co-localisation of SYCE2 and RAD51 foci was dependent on SYCP1. SIM super-resolution imaging of asynapsed pachytene *Sycp1*^*-/-*^ spermatocytes showed that SYCE2 microfoci still frequently co-localised with RAD51 foci (Figure 4C), and quantitative analysis confirmed the strong enrichment of co-localised SYCE2-RAD51 foci, with centroids separated by approximately 80 nm (Figure 4D). Interestingly, the co-localised SYCE2-RAD51 centroid distances and RAD51 foci frequency of *Sycp1*^*-/-*^ spermatocytes were more similar to *Syce3*^*WY/WY*^ than *Syce3*^*Δ/Δ*^ spermatocytes (Figure 4B, Figure 4D, Supplementary Figure S2B). Thus, SYCP1 is not required for the association between SYCE2 microfoci and RAD51 foci, but SYCP1 tetramer lattices may directly or indirectly influence the structure and/or progression of recombination intermediates.

In *Syce3* and *Sycp1* asynapsed pachytene spermatocytes, most autosomal axes were unambiguously paired with axes of similar length, but remained separated by distances greater than the ∼200 nm distance separating SYCP3 axis traces in fully synapsed wild-type pachytene nuclei (Supplementary Figure S3A). The ∼200 nm distance between traces along the centre of each SYCP3 immunstained axis in chromosome spreads is as expected larger than the ∼100 nm gap between lateral elements measured by electron microscopy of fixed cells (Westergaard and von Wettstein, 1972). We noticed that SYCE2 microfoci and co-localised SYCE2-RAD51 foci were frequently found between paired axes, at ∼80-120 nm from the nearest SYCP3 axis trace, located where the CE assembles in the mature SC (Figure 4E; Supplementary Figure S3B; Supplementary Figure S3C). This was also the case in *Sycp1*^*-/-*^ spermatocytes, which lack transverse filaments (Figure 4E). Thus, co-localised SYCE2-RAD51 foci are enriched in regions between paired chromosomes, where the central element would normally assemble, independently of SYCP1. Hence, its localisation is likely driven by the underlying architecture of DNA and/or recombination intermediates at these sites.

### Subsets of SYCP1 foci and SYCE2 microfoci are associated with sites of potential synapsis in *Syce3* and *Sycp1* mutant spermatocytes

Paired axes in asynapsed pachytene nuclei are typically separated by >200 nm, but contain regions of potential synapsis where axis are separated by ∼200 nm. Hence, we wondered whether SYCP1 foci and/or SYCE2 microfoci might be associated with regions of close axis proximity and attempted synapsis within asynapsed pachytene *Syce3* and *Sycp1* mutant spermatocytes.

Given the difference in size and frequency of SYCP1 foci (Crichton et al., 2022)(Supplementary Figure S1B), we expected that any stabilising effect of SYCP1 on synapsis would result in closer axis proximity in *Syce3*^*Δ/Δ*^ than *Syce3*^*WY/WY*^ spermatocytes. However, most SYCP1 foci were not associated with regions of close axis proximity in *Syce3*^*Δ/Δ*^ nuclei and only a subset corresponding to the largest SYCP1 foci were enriched at these sites (Figure 5A). In contrast, large SYCP1 foci were not associated with close axis proximity in *Syce3*^*WY/WY*^ nuclei (Figure 5A). Thus, large SYCP1 foci, likely corresponding to SYCP1 tetramer lattices, are enriched at sites of attempted synapsis in *Syce3*^*Δ/Δ*^ spermatocytes.

**Figure 5.**
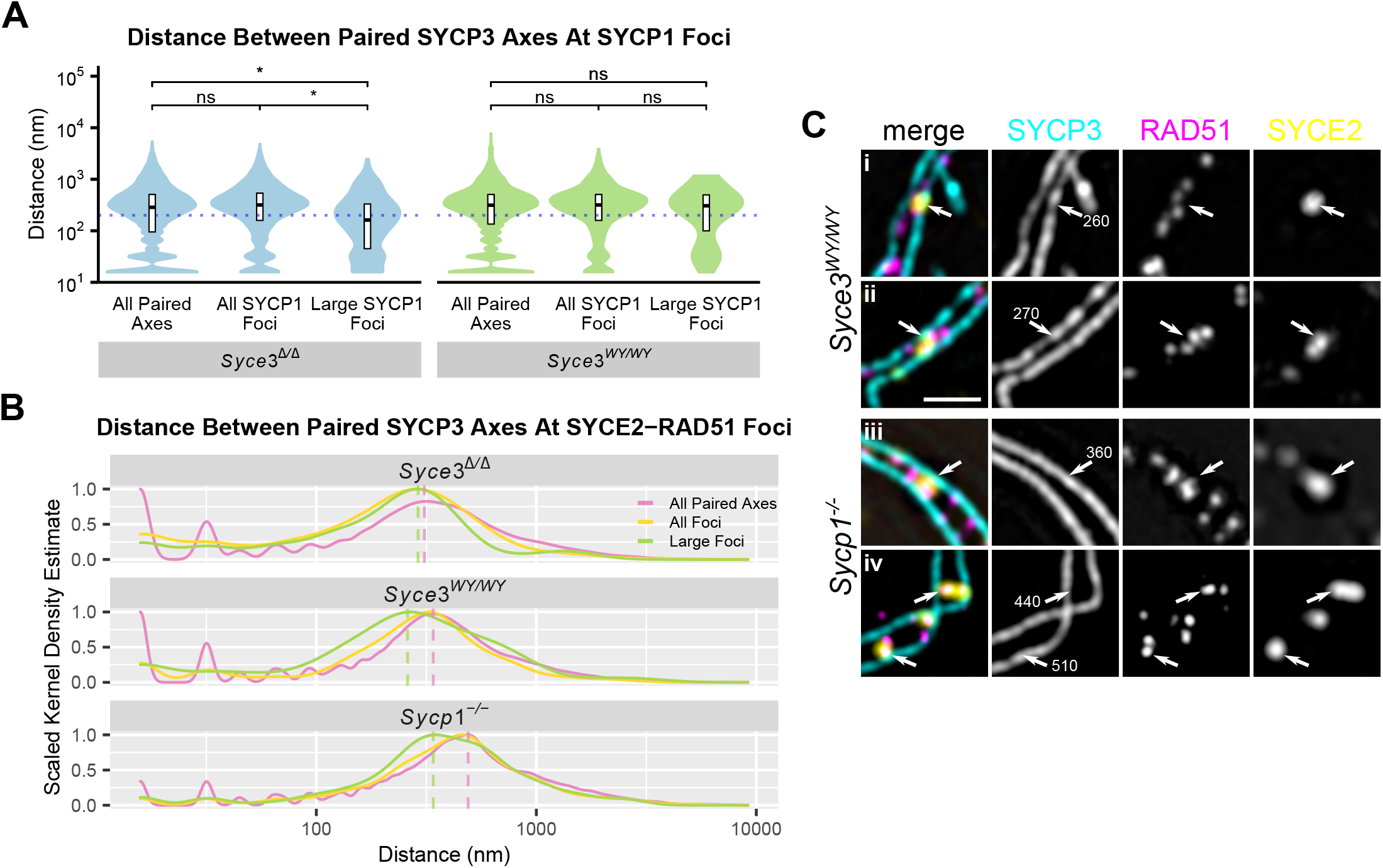
SYCP1 foci and SYCE2-RAD51 foci are enriched for regions of close axis proximity in *Syce3* and *Sycp1* mutant spermatocytes. (**A**) The nearest distance between paired SYCP3 axes traces in asynapsed pachytene *Syce3* nuclei was determined for each point on the SYCP3 axes trace and for those points closest to all or large (> 150 px^2^) axial SYCP1 foci (Supplementary Figure S1B). Crossbars represent quartiles; blue dotted line indicates 200 nm; *, p< 0.01; ns, not significant (Mann-Whitney U test, medians are 272.5, 303.6, 158.5, 317, 317 and 302.2 nm, n=24, 24, 24, 23, 23, 12 nuclei); 3 animals analysed for each *Syce3* genotype. Summary statistics are in Supplementary Table ST2. (**B**) Kernel density estimates that paired SYCP3 axis traces are separated by the indicated distance at all or large axial SYCE2-RAD51 foci. The 10% SYCE2 microfoci with the largest areas within each genotype were classed as large. Dotted lines represent approximate positions of the peak populations in *Syce3*^*Δ/Δ*^, *Syce3*^*WY/WY*^ and *Sycp1*^*-/-*^ asynapsed pachytene spermatocytes (310, 340 and 450 nm for all paired axes; 290, 260 and 340 nm for large foci); 3 *Syce3*^*Δ/Δ*^, 3 *Syce3*^*WY/WY*^ and 2 *Sycp1*^*-/-*^ animals analysed. (**C**) SIM super-resolution images of paired SYCP3 regions representing peak populations associated with large SYCE2 microfoci in (**B**). Chromosome spreads were immunostained for SYCP3 (blue), RAD51 (magenta) and SYCE2 (yellow). Axial SYCE2-RAD51 co-foci are indicated with arrows, numbers in the SYCP3 panel indicate the axis separation (nm). Scale bar, 1 µm.

In contrast, in *Syce3*^*WY/WY*^ and *Sycp1*^*-/-*^ spermatocytes, large SYCE2 microfoci that co-localised with RAD51 foci were strongly enriched at regions of intermediate axis proximity (Figure 5B). Axis separation in these regions (∼260 nm and ∼340 nm in *Syce3*^*WY/WY*^ and *Sycp1*^*-/-*^ spermatocytes, respectively) was not as close as the ∼200 nm separation in full synapsis and at large SYCP1 foci in *Syce3*^*Δ/Δ*^ spermatocytes (Figure 5A, Supplementary Figure S3A). Thus, in mutants where assembly of SYCP1 tetramer lattices and integrated SYCP1-SYCE3 lattices is perturbed, a subset of co-localised SYCE2-RAD51 foci are associated with sites of intermediate axis proximity that could represent pre-synaptic interactions between paired axes. SIM super-resolution imaging revealed that these sites typically consisted of single large SYCE2 microfoci overlapping multiple RAD51 foci, and bridging gaps of ∼260 nm and up to ∼500 nm between paired SYCP3 axes in *Syce3*^*WY/WY*^ and *Sycp1*^*-/-*^ spermatocytes, respectively (Figure 5C). Thus, large SYCE2-RAD51 foci appear to represent bridges of SYCE2 physically linking paired pre-synaptic axes at RAD51 recombination sites.

Taken together with existing data in the field, our analyses suggest that SYCE2 can be recruited to RAD51 recombination sites, form physical bridges between paired pre-synaptic chromosome axes at these sites, and localise to central regions between paired chromosome axes, independently of SYCP1. Further, SYCP1 lattice assembly promotes initiation and/or stabilisation of synaptic interactions between paired chromosome axes, independently of SYCE2. Hence, we propose that these two pathways act in parallel to recruit SC components to meiotic chromosomes in mouse meiosis.

### The recombinant SYCE2-TEX12 complex interacts with SYCP1 and DNA *in vitro*

Our *in vivo* analyses suggest that there may be as yet uncharacterised direct interactions between SYCE2 and recombination foci (Figure 4) that have not yet been detected biochemically. Furthermore, given the focal rather than filamentous appearance of SYCP1-independent SYCE2, we reasoned that the rare co-localisation between SYCP1 and SYCE2 that we detected in *Syce3*^*Δ/Δ*^ spermatocytes (Figure 3D) might represent SYCE3-independent interactions between SYCP1 and SYCE2 that also have not yet been detected biochemically. We therefore tested for these interactions *in vitro* using purified bacterially-expressed recombinant proteins. We identified biochemically that SYCP1 interacts with SYCE2-TEX12, but not SYCE1-SIX6OS1, through SYCP1’s unstructured N-terminus (Figure 6A, Figure 6B, Supplementary Figure S4). This interaction is low-affinity and does not alter SYCP1 structure (Figure 6C, Figure 6D), so is likely stabilised by cooperativity within an SC lattice. Further, recombinant SYCE2-TEX12 interacts with single- and double-stranded DNA *in vitro* (Figure 6E), so possesses the ability to bind DNA species present at recombination sites. In combination with existing data showing that SYCP1 can also bind to DNA, and that SYCE3 interacts with SYCP1, SYCE1-SIX6OS1 and SYCE2-TEX12, these interactions rationalise our *in vivo* data, and suggest an important symmetry in which SYCP1 and SYCE2-TEX12 bind to DNA and each other, self-assemble into architectural SC components, and can independently initiate SC assembly (Figure 6F).

**Figure 6.**
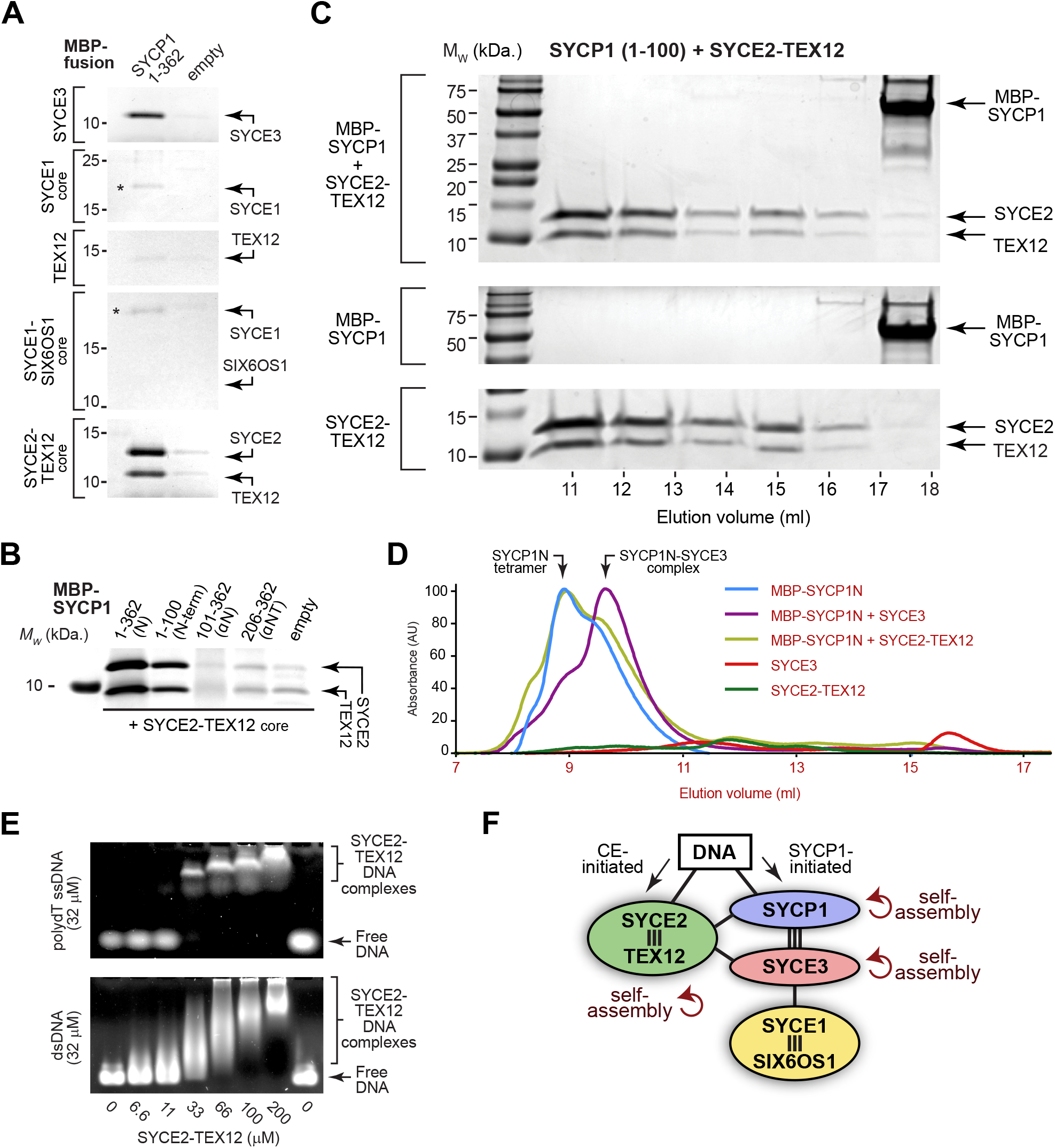
SYCE2-TEX12 biochemically interacts with both SYCP1 and DNA *in vitro*. (**A**) Amylose pull-down of central element proteins and complexes following incubation with MBP-SYCP1N (amino-acids 1-362) and free MBP; a degradation product of MBP-SYCP1N is indicated by an asterisk. Uncropped gels are shown in Supplementary Figure S4. (**B**) Amylose pull-down of SYCE2-TEX12 following recombinant co-expression with MBP-SYCP1 constructs and free MBP (empty). Uncropped gels are shown in Supplementary Figure S4. (**C**) Size-exclusion chromatography of MBP-SYCP1 amino-acids 1-100 alone and upon equimolar incubation with SYCE2-TEX12, shown as SCS-PAGE of elution fractions. (**D**) Size-exclusion chromatography of MBP-SYCP1N (amino-acids 1-362) alone and upon equimolar incubation with SYCE3 and SYCE2-TEX12, along with SYCE3 and SYCE2-TEX12 alone, shown as UV absorbance (280 nm) chromatograms. Given their differences in amino-acid composition, absorbance is dominated by MBP-SYCP1N, with SYCE3 and SYCE2-TEX12 absorbing weakly, so chromatograms largely indicate the elution profiles of MBP-SYCP1N species. (**E**) Electrophoretic mobility shift assays analysing the ability of purified SYCE2-TEX12 protein complexes to interact with 32 µM (per base) polydT single-stranded DNA (ssDNA, top) and double-stranded DNA (dsDNA, bottom) substrates *in vitro*. (**F**) Interaction network of SC central element proteins, indicating strong interactions (three lines), weak interactions (single lines) and self-assembly characterised biochemically with purified proteins *in vitro* (modified from Crichton et al., 2022).

### Integration of parallel SYCP1- and CE-initiated pathways of SC assembly *in vivo*

Do the parallel pathways recruiting SC components in *Syce3* and *Sycp1* mutant spermatocytes while they are arrested in pachytene reflect pathways operating normally during SC assembly in wild-type meiosis? SIM super-resolution imaging detected SYCE2 microfoci on unsynapsed regions of SYCP3 axes, frequently co-localised with RAD51 foci, during leptotene in *Sycp1*^*+/-*^ and *Sycp1*^*-/-*^ spermatocytes, suggesting that SYCE2 is recruited to recombination foci during early stages of SC assembly that precede asynapsis and pachytene arrest (Supplementary Figure S5A, Supplementary Figure S5B). Further, in leptotene and zygotene wild-type spermatocytes from C57BL/6J mice, SoRa super-resolution imaging detected independent SYCP1 foci and SYCE2 microfoci on unsynapsed regions of meiotic chromosome axes (Figure 7A). We also detected co-localised SYCE2-SYCP1 foci on unsynapsed axes in wild-type spermatocytes (Figure 7A). Hence, independent recruitment of SYCE2 and SYCP1 to chromosome axes, in addition to some SYCE2-SYCP1 co-localisation, all occur prior to synapsis in wild-type meiosis.

**Figure 7.**
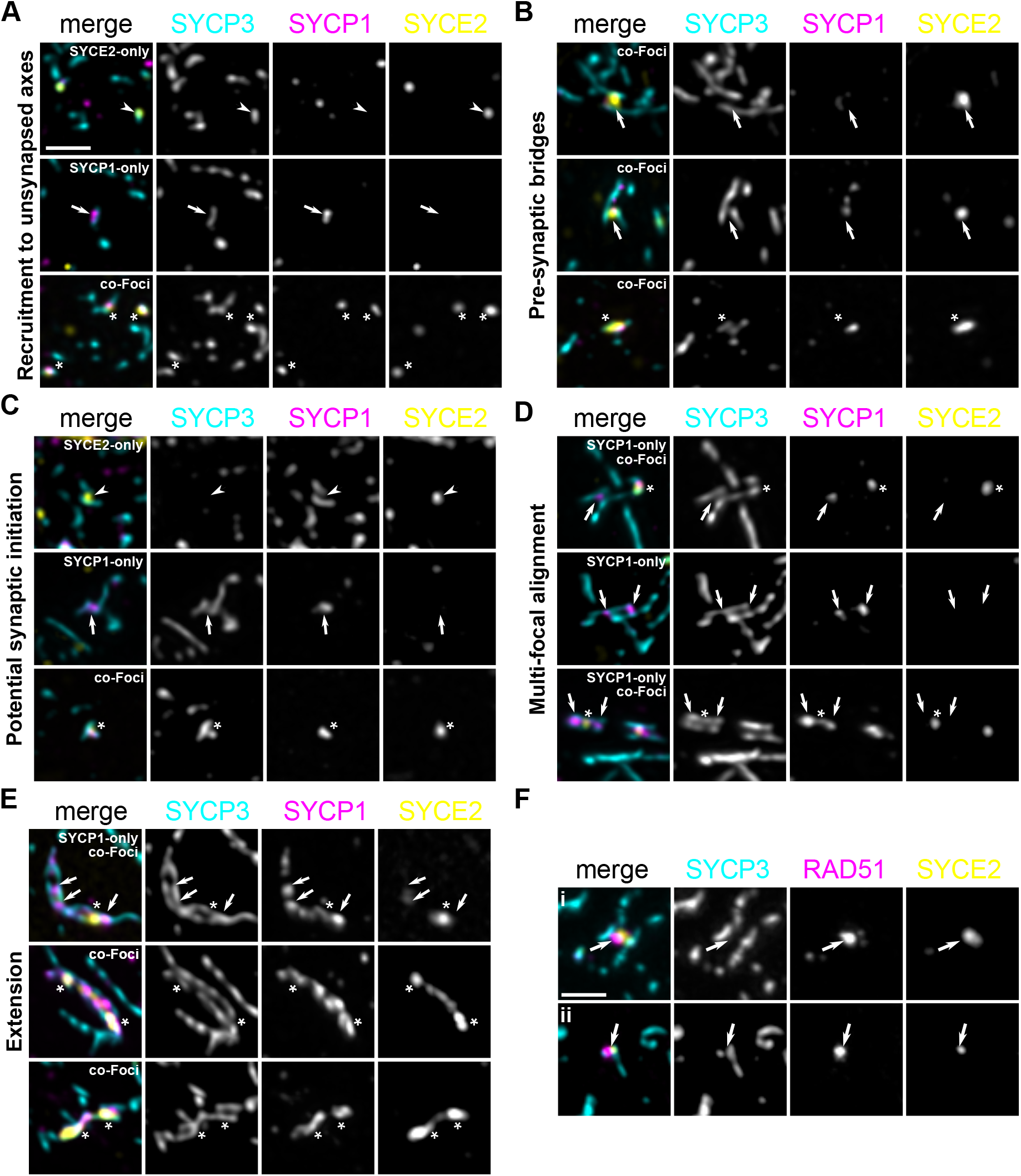
Synaptonemal complex assembly in wild-type spermatocytes. (**A**-**E**) SoRa images of C57BL/6J leptotene spermatocyte nuclei immunostained for SYCP3 (blue), SYCP1 (magenta) and SYCE2 (yellow). Foci are highlighted with arrowheads (SYCE2-only), arrows (SYCP1-only) or asterisks (SYCE2-SYCP1 co-localised foci) are indicated. Scale bar, 1 µm. (**A**) Unsynapsed SYCP3 axes with SYCE2, SYCP1 or SYCE2-SYCP1 co-localised foci. (**B**) SYCP3 axes separated by >300 nm and <500nm, and physically linked by SYCE2. (**C**) SYCP3 axes separated by ∼200 nm and physically linked by SYCE2, SYCP1 or SYCE2-SYCP1 co-localised foci. (**D**) Longer regions of SYCP3 axes, separated by ∼200 nm and physically linked by multiple SYCE2, SYCP1 or SYCE2-SYCP1 co-localised foci. (**E**) As (**D**) but with linear extensions of SYCE2 or SYCP1 extending out from foci along the length of the SYCP3 axis. (**F**) SoRa images of C57BL/6J leptotene spermatocyte nuclei immunostained for SYCP3 (blue), RAD51 (magenta) and SYCE2 (yellow). SYCE2-RAD51 co-localised foci are highlighted with arrows. Axis separation in (i) is in the ∼300-500 nm range described for (**B**). Scale bar, 1 µm.

We also detected SC assembly intermediates in wild-type leptotene and zygotene spermatocytes that resembled the structures detected at regions of intermediate axis proximity in *Syce3*^*WY/WY*^ and *Sycp1*^*-/-*^ asynapsed pachytene spermatocytes. These large pre-synaptic bridges of SYCE2 connected SYCP3 axes separated by 300-500 nm and co-localised with RAD51 (Figure 7B, Figure 7F). We also observed regions in wild-type spermatocytes where axes were separated by ∼200 nm, approximately the same distance as full synapsis, and physically connected by separate or co-localised SYCP1 foci and SYCE2 microfoci (Figure 7C). Hence, SYCE2 pre-synaptic bridges, their co-localisation with RAD51 sites, and the association of SYCP1 with regions of synapsis-like axis proximity, were similarly observed in *Syce3* and *Sycp1* spermatocytes arrested in pachytene, and in wild-type spermatocytes progressing through leptotene and zygotene.

Finally, we observed unique intermediate structures in wild-type spermatocytes that were not present in *Syce3* and *Sycp1* mutants. Specifically, we identified long regions of synapsing axes in which multiple SYCP1 foci or SYCP1-SYCE2 co-foci simultaneously contacted both axes (Figure 7D). In some cases, SYCP1 or SYCE2 staining extended linearly away from these foci, along chromosome axes, to link and coalesce adjacent foci (Figure 7E). These SYCP1 and SYCE2 extensions may correspond to the growth of SYCP1-containing lattices and SYCE2-TEX12 fibres, respectively (Dunce et al., 2021, 2018). The formation of these self-assembling structures is promoted by SYCE3 *in vitro* (Crichton et al., 2022), explaining their absence in *Syce3* and *Sycp1* mutants.

Overall, our findings are consistent with parallel pathways recruiting SC proteins to meiotic chromosomes, generating multiple independent nucleation sites that extend and coalesce, to drive assembly of the mammalian SC.

## Discussion

### Linking SC assembly to recombination sites in mammals

The recruitment of SYCE2 to recombination sites in pre-synaptic bridges and synaptic initiation sites establishes a hitherto elusive link between recombination and SC initiation in mammals. This novel pathway of mammalian SC assembly provides an important link with *Sordaria* and budding yeast, in which SC initiation is known to occur at recombination sites (Cahoon and Hawley, 2016; Gao and Colaiácovo, 2018; Zickler and Kleckner, 2015). SYCE2 recruitment to recombination sites is consistent with previous findings that SYCE2 co-immunoprecipitates with RAD51 (Bolcun-Filas et al., 2009) and, given the binding between SYCE2-TEX12 and both dsDNA and ssDNA *in vitro*, may occur through direct interactions. It may also be facilitated by an adapter protein, such as TEX11, whose orthologue Zip4 has been implicated in the association of analogous CE complex Gmc2-Ecm11 with recombination and synapsis initiation sites in budding yeast (Pyatnitskaya et al., 2022).

The pre-synaptic bridges observed in leptotene likely correspond to RAD51/DMC1-containing connections between axes (Tarsounas et al., 1999), and potentially to pre-synaptic bridges seen in other species (Zickler and Kleckner, 2015). Hence, SYCE2 localisation to these sites, independently of SYCP1, suggests that CE components play much earlier roles in SC assembly than previously thought. SC initiation by SYCE2 could be regulated and/or restricted to specific recombination intermediates, explaining why only a subset of SYCE2-RAD51 foci is associated with synapsis initiation in *Syce3*^*WY/WY*^ mice. However, it is not clear whether, like in *Sordaria* and budding yeast (Fung et al., 2004; Zhang et al., 2014), recombination sites associated with SC initiation are fated to become genetic crossovers.

### Parallel recruitment pathways in the mammalian SC

Our data, in combination with previous data in the field, suggest that the mammalian SC assembles using parallel recruitment pathways: one pathway that recruits SYCP1 transverse filaments to synapsing chromosome axes (Figure 8Ai), and a separate pathway that recruits the CE proteins SYCE2-TEX12 to recombining DNA on pre-synaptic chromosome axes (Figure 8Aii). Subsequent extension and coalescence of independently-nucleated assemblies then allow the SC to extend along the length of the chromosomes (Figure 8B). The co-existence of parallel recruitment pathways is consistent with the complex relationship between later stages of SC assembly and recombination in mammals (Gruhn et al., 2016), and may safeguard SC assembly in a chromosomal landscape where the recombination machinery and genome-wide distribution of recombination is evolving rapidly (Baudat et al., 2013).

**Figure 8.**
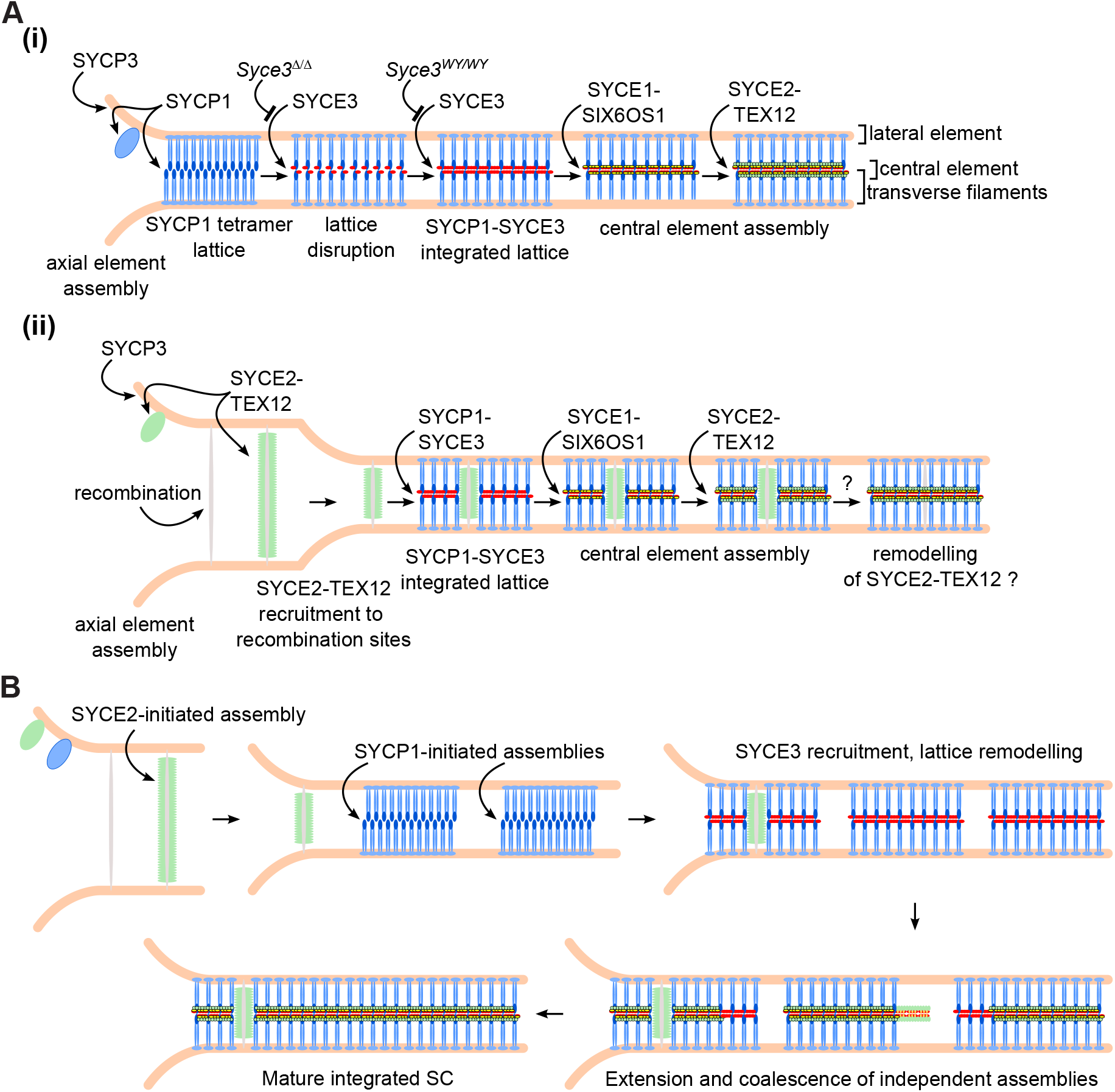
Model for mammalian synaptonemal complex assembly (**A**) Parallel recruitment pathways contributing to mammalian SC assembly (**i**) SYCP1-initiated assemblies recruit SYCP1 to unsynapsed axes and form SYCP1 tetramer lattices between chromosome axes in close ∼200 nm proximity. Recruitment of SYCE3 disrupts and remodels the SYCP1 tetramer lattice, and SYCE3 self-assembly interactions drive extension of the SYCP1-SYCE3 integrated lattice. SYCE3 recruits the remaining central element components SYCE1-SIX6OS1 and SYCE2-TEX12 to stabilise the extending lattice. (**ii**) Potential pathway for SYCE2-initiated SC assembly. Recombining DNA directly recruits SYCE2-TEX12 to unsynapsed axes and inter-homolog recombination sites. SYCE2 assemblies bridge pre-synaptic paired chromosome axes separated by up to ∼500 nm. Progression of synapsis and interactions between SYCE2-TEX12 and SYCP1-SYCE3 allows association with integrated SYCP1-SYCE3 lattices in regions where axes are separated by ∼200 nm. Recruitment of SYCE1-SIX6OS1 and SYCE2-TEX12 allows the central element to assemble. It is not clear whether remnants of the SYCE2-TEX12 pre-synaptic bridge assembly persist as a structural heterogeneity at recombination sites in the mature SC, or are remodelled by the SC extending into these regions. (**B**) Integration of parallel recruitment pathways for SC assembly. SYCP1-independent recruitment of SYCE2-TEX12 to recombination sites and SYCE2-independent recruitment of SYCP1 occurs on unsynapsed axes from leptotene. SYCE2-initiated assemblies can form bridges between pre-synaptic axes separated by up to ∼500 nm. As synapsis bring axes into ∼200 nm proximity, SYCP1-initiated lattice assemblies can form links between axes. Recruitment of SYCP1-SYCE3 to SYCE2-initiated assemblies, and SYCE3-dependent remodelling of SYCP1-initiated assemblies promotes formation of integrated SYCP1-SYCE3 lattices. Polymerisation of SYCE2-TEX12 filaments and SYCP1-SYCE3 integrated lattices drives extension of independently-nucleated assemblies, which coalesce to form the mature integrated SC. As in (A), it is not clear whether remnants of the SYCE2-TEX12 pre-synaptic bridge assemblies persist as at recombination sites in the mature SC, or are remodelled. Note that although the SYCP1-independent and SYCE2-independent pathways are genetically separable, in wild-type spermatocytes early foci and assemblies may contain both SYCE2-TEX12 and SYCP1 (co-localised foci in Fig7), likely in combination with SYCE3.

SYCP1-initiated SC assembly (Figure 8Ai) is likely instigated by the formation of SYCP1 tetramer lattices on unsynapsed chromosome axes – possibly linking sister chromatids rather than homologs, bridging between loops of an individual chromatid, or as unbridged assemblies. SYCP1 tetramer lattices then establish inter-homologue links by SYCP1 molecules on one side of the lattice capturing the homolog axis at the necessary ∼200 nm proximity, and achieving the necessary cooperativity to stabilise these inter-homolog linkages. The majority of SYCP1 tetramer lattices are then remodelled by SYCE3 into integrated SYCP1-SYCE3 lattices, enabling the recruitment of CE complexes SYCE1-SIX6OS1 and SYCE2-TEX12, which confer stability to the growing SC structure (Dunce et al., 2018) (Crichton et al., 2022).

In contrast, the mechanism of CE-initiated SC assembly remains uncertain (Figure 8Aii). SYCE2 could recruit and nucleate SYCP1 tetramer lattice formation through the direct interaction between SYCP1 and SYCE2-TEX12 reported here, potentially through non-CE-like structures similar to that shown in Figure 3D. However, the limited association between SYCE2 foci and SYCP1 foci in *Syce3*^*Δ/Δ*^ spermatocytes suggests that any recruiting interactions between these proteins depend on SYCE3. Hence, SYCE2-TEX12 may directly recruit and nucleate assembly of integrated SYCP1-SYCE3 lattices at recombination sites. It is uncertain whether SYCE2 at recombination sites adopts the same fibrous SYCE2-TEX12 assemblies that are formed within the CE (Dunce et al., 2021). The difference in staining intensity between SYCE2 microfoci and the CE means that it is not possible to determine whether SYCE2 structures at recombination sites remain intact upon full SC assembly. Nevertheless, it is intriguing to speculate that SYCE2 bound to recombining DNA could create a channel or irregularity in the integrated SYCP1-SYCE3 lattice that might facilitate recombination. Indeed, such a mechanism could explain the structural differences observed within the SC at recombination sites in *C. elegans* (Libuda et al., 2013).

### Interaction symmetry within the SC

The SC’s interaction network includes an intriguing symmetry in which SYCP1 and SYCE2-TEX12 both bind to DNA, to each other, self-assemble into architectural components of the SC, and initiate SC assembly (Figure 6F). Thus, SYCP1 and SYCE2-TEX12 appear to be the most fundamental components of the SC, in agreement with their early evolutionary origin (Fraune et al., 2016). Hence, it is possible that SYCE2’s early evolutionarily origin is attributed to its role in recombination, with its function in CE assembly having evolved later, after the emergence of SYCE1 and SIX6OS1. The presence of multiple reinforcing and self-assembly interactions within the SC could allow different SC components to nucleate and drive assembly of this structure in different contexts or in different species. Thus, it is possible that SC assembly is more varied and heterogeneous than the two pathways described herein.

The DNA-binding abilities of SYCP1 and SYCE2-TEX12 could relate to their recruitment to chromosome axes, or recombination sites, or both. In SYCP1, DNA-binding is mediated by its C-terminus, located with the SC’s lateral element, so is primarily attributed to its chromosomal recruitment (Dunce et al., 2018). Nevertheless, it is possible that its multiple DNA-binding sites have additional roles in stabilising and/or regulating recombination intermediates. Hence, SYCP1 lattices may directly or indirectly influence recombination, explaining the differences in RAD51 foci frequency of *Sycp1*^*-/-*^ and *Syce3*^*WY/WY*^ mutants in comparison with *Syce3*^*Δ/Δ*^ spermatocytes (Figure 4, Supplementary Figure S2). The localisation of SYCE2-TEX12 to the midline between paired axes in pre-synaptic microfoci and in *Sycp1*^*-/-*^ spermatocytes suggests that these CE components can be recruited and positioned by the underlying structure of DNA, likely in the form of recombination intermediates. SYCE2-TEX12 binding could stabilise and/or regulate these recombination intermediates, and could allow SYCE2 to impact on DNA repair in non-meiotic contexts (Hosoya et al., 2018). These observations point to a role for the underlying structure of DNA at recombination sites in driving evolution of their support proteins, including the establishment of transverse filament coiled-coil proteins of the correct length to span between recombining axes. Hence, the underlying DNA may have provided the architectural specifications to which the proteinaceous scaffold has evolved.

## Materials and Methods

### Recombinant protein expression and purification

SYCP1, SYCE3, SYCE1, SYCE1-SIX6OS1, TEX12 and SYCE2-TEX12 protein constructs and complexes were purified as previously described (Davies et al., 2012; Dunce et al., 2021, 2018; Dunne and Davies, 2019a, 2019b; Sánchez-Sáez et al., 2020). In general, proteins were expressed as His_6_- or His_6_-MBP fusions in BL21(DE3) *E. coli* cells (Novagen^®^), and purified from lysate through Ni-NTA (Qiagen) or amylose (NEB) affinity, with removal of the tag by TEV protease treatment, followed by anion exchange chromatography (HiTrap Q HP, Cytiva) and size exclusion chromatography (HiLoad™ 16/600 Superdex™ 200, Cytiva) in 20 mM Tris pH 8.0, 150 mM KCl, 2 mM DTT. Purified proteins were concentrated using Amicon Ultra^®^ 10,000 MWCO centrifugal filter units (Millipore) or Microsep™ Advance 3kDa (PALL) centrifugal filter units, and flash-frozen in liquid nitrogen for storage at -80°C. Samples were analysed by Coomassie-stained SDS-PAGE, and concentrations were determined using a Cary 60 UV spectrophotometer (Agilent) with molecular weights and extinction coefficients calculated by ExPASY ProtParam (http://web.expasy.org/protparam/).

### Pull-down assays

To determine central element interactions of SYCP1N (amino-acids 1-362), 100 μl of MBP-SYCP1N at 2 mg/ml was incubated for 20 minutes on ice with 100 ul amylose resin slurry (NEB) equilibrated in 20 mM Tris pH 8.0, 50 mM KCl, 2 mM DTT, 0.1% Tween-20. After centrifugation at 3500 rpm for 1.5 minutes, the supernatant was removed and 500 μl of 20 mM Tris pH 8.0, 150 mM KCl, 2 mM DTT, 0.1% Tween-205 mg/ml BSA was added and incubated for 20 minutes on ice. After centrifugation at 3500 rpm for 1.5 minutes and removal of the supernatant, a central element protein, or complex thereof, was applied at a three-fold molar excess with respect to MBP-SYCP1αN and incubated for 20 minutes at 37 °C. The mixture was centrifuged at 3500 rpm for 1.5 minutes, and the supernatant removed. The resin was washed thrice with 20 mM Tris pH 8.0, 50 mM KCl, 2 mM DTT, 0.1% Tween-20, once with 20 mM Tris pH 8.0, 500 mM KCl, 2 mM DTT, 0.1% Tween-20, and after removal of the supernatant, 25 μl loading buffer and 4 μl beta-mercaptoethanol was added and heated at 95 °C prior to analysis by SDS-PAGE.

### Gel filtration assays

50 µl protein samples were prepared corresponding to MBP-SYCP1N, SYCE3, SYCE2-TEX12, and 1:1 mixtures of MBP-SYCP1N with SYCE3 and with SYCE2-TEX12, in which each component was present at 235 µM. Samples were incubated for 1 hour at room temperature and centrifuged at 14000 g at 4°C for 30 minutes. Size exclusion chromatography was performed using a Superdex™ 200 Increase 10/300 GL column in 20 mM Tris pH 8.0, 150 mM KCl, 2 mM DTT at 0.5 ml/min. Chromatograms correspond to absorbance at 280 nm.

### Electrophoretic mobility shift assay

polydT ssDNA or dsDNA at 33 μM per nucleotide or base pair was incubated with the indicated concentrations of protein for 1 hour in 50 mM triethanolamine pH 7.5, 2 mM DTT, and resolved on a 0.5% w/v agarose gel in 1x TAE buffer in the cold room for 5 hours. DNA was detected by SYBR™ Gold (ThermoFisher) staining using a Typhoon™ FLA 9500 (GE Healthcare), with 473 nm laser at excitation wavelength 490 nm and emission wavelength 520 nm, using the LPB filter and a PMT voltage of 400 V.

### Mice and Meiotic Chromosome Spreads

Meiotic chromosome spreads from the testes of 2-4 months old F0 founder *Syce3* mutant mice generated by CRISPR/Cas9 gene editing in zygotes were as described (Crichton et al., 2022). Meiotic chromosome spreads from same adult F0 *Syce3* animals that were characterized previously (Crichton et al., 2022) were used in this study. No regions of phenotypic mosaicism were detected in the testes of these F0 animals (Crichton et al., 2022). Adult C57BL/6J animals (Charles River) were culled by cervical dislocation at 2-4 months old, and their testes used to make meiotic chromosome spreads (Costa et al., 2005). Chromosome spreads from *Sycp1*^*-/-*^ mice were prepared as described (Slotman et al., 2020). Chromosome spreads were stained with antibodies (Supplementary Table ST1) as described (Crichton et al., 2017) and mounted under high precision coverslips (Marienfeld) with antifade mounting medium (Vectashield).

Spermatocytes were staged base on SYCP3 staining of chromosome axes. Leptotene spermatocytes had short, fragmented filaments of axial staining. Zygotene axes were extended, but with limited alignment of chromosome pairs. Pachytene spermatocytes had complete and close alignment of autosomes, only possible to resolve using super-resolution techniques. Asynapsed pachytene spermatocytes from mutant lines had complete axis formation and extensive loose alignment of autosomes, sufficiently distant to resolve by diffraction-limited widefield microscopy.

### Widefield Fluorescent Imaging

Single plane images (SYCP3 + TEX12, SYCE1, or SYCE3) were captured using a Zeiss Axioplan II fluorescence microscope equipped with a Photometrics Coolsnap HQ2 CCD camera, while nuclei captured across multiple z-planes (SYCP3 + SYCE2) were imaged using a Photometrics Prime BSI CMOS camera coupled to a Zeiss AxioImager M2 fluorescence microscope. Micromanager (Version 1.4) and Huygens Essential were used to capture images and deconvolve z-stacks respectively, and maximum intensity projections and analysis performed using Fiji (Schindelin et al., 2012).

### Super-Resolution Imaging

3D SIM images were captured using a Nikon N-SIM microscope equipped with an Andor iXon 897 EMCCD camera (Andor technologies, Belfast UK). 3D SoRA images were captured using a Nikon Eclipse Ti2 microscope with a CSU-W1 SoRa unit (Yokogawa) and a Photometrics Prime 95B CMOS camera and deconvolved with Nikon NIS-Elements. Capture parameters were consistent for each antibody combination. Nuclei larger than the field of view were captured as tiled images with 15% overlap (Supplementary Figure S6), and stitched together in Nikon NIS-Elements (SIM) by blending, or in Fiji by pairwise stitching in (SoRa) (Preibisch et al., 2009). Images were maximum intensity projected for further analysis. Quantitative analysis was performed using custom pipelines in Fiji, Python and R.

### Quantitative Image Analysis

Binary masks were generated in Fiji by manual thresholding of antibody staining, and drawing a region of interest around DAPI staining for nuclear territories. Masks of focal staining patterns were converted into labelmaps for further analysis. SYCE2 masks from synapsed pachytene control images were reduced to a single pixel line using the Skeletonize function. SYCP3 axis skeletons were generated by tracing with the Simple Neurite Tracer plugin, skeletons orientated by bright centromeric DAPI signal, and results saved in the .traces format. Axes typically showed alignment with a single partner of comparative size. These unambiguous pairings were manually identified and recorded within the .traces file. Only autosomal axes with an identified partner, which could be traced for their full length and orientated were included. Further analysis was performed using Python3 and R.

The traces which comprise SYCP3 axis skeletons were read into Python using existing code (https://github.com/huangziwei/TracePy). Distances from each pixel of a SYCP3 skeleton to their previously identified partner were calculated by querying a k-d tree of coordinate values generated for this target. Distances from SYCE2 skeleton coordinates and weighted centroids of foci to the SYCP3 mask, SYCP3 skeletons, or to weighted centroids of other foci were all calculated by the same method. Orientation of foci towards or away from their partner axis was calculated by comparing the distance between paired axes at that location, and the distance from the focus to the same point on the partner axis. Focus labels from different channels sharing any coordinates were considered to be overlapping. Foci less than 3 square pixels in area were excluded from all analysis as background.

Foci were shuffled within the nucleus by randomly assigning their labels with new centroid coordinates within the nuclear territory, ensuring edges of each label did not overlap each other or extend beyond the nuclear mask. For shuffling within the axes, foci whose centroids fell within the SYCP3 mask territory were similarly shuffled within the SYCP3 mask territory, with the exception that the focus edge was permitted to exceed the boundary of the SYCP3 mask, as was often the seen in the original images. For SYCP1 foci shuffling, only foci greater than 3 square pixels in area were used for shuffling. For SYCE2 microfoci shuffling, only foci greater than 3 square pixels and less than 100 square pixels in area were used for shuffling. When comparing observed data and shuffled data only foci used in shuffling were included in the observed data.

## Animal Ethics

Institutional and national ethical and welfare guidelines and regulations were followed for all experiments involving animals. Experiments were approved by the University of Edinburgh animal welfare and ethics review board and were performed under UK Home Office licences PP3007F29 and PB0DC8431.

## Acknowledgements

We are grateful to A. Wheeler, L. Murphy and M. Pearson in the IGC Advanced Imaging Resource for their support with imaging and image analysis, the Edinburgh Super-Resolution Imaging Consortium for super-resolution imaging, and the University of Edinburgh Bioresearch & Veterinary Services for mouse husbandry. We thank Howard Cooke and Ricardo Benavente for generously sharing reagents. We thank Wendy Bickmore, Javier Caceres, Cova Vara and Adele Marston for critically reviewing the manuscript. I.R.A. and J.H.C. are supported by MRC University Unit grant MC_UU_00007/6. O.R.D. is a Wellcome Senior Research Fellow (Grant Number 219413/Z/19/Z).

## Author contributions

J.H.C. prepared and imaged chromosome spreads, developed imaging analysis pipelines and analysed imaging data. J.M.D. performed biochemical and biophysical experiments. W.M.B. provided materials and contributed to editing the manuscript. O.R.D. and I.R.A. analysed data, designed experiments and wrote the manuscript.

