## SupplementaryMaterial for "Parallel recruitment pathways contribute to synaptonemal complex assembly during mammalian meiosis"

#### Supplementary Tables

| Antibody | Dilution | Source |
|---|---|---|
| Mouse anti-SYCP3 | 1:500 | Abcam #ab97672 |
| Rabbit anti-SYCP1 | 1:200 | Abcam #ab15090 |
| Rabbit pig anti-SYCE1 | 1:200 | Howard Cooke (MRC HGU, Edinburgh)<br>(Costa et al., 2005) |
| Guinea pig anti-SYCE2 | 1:1000 | Howard Cooke (MRC HGU, Edinburgh)<br>(Bolcun-Filas et al., 2009) |
| Guinea pig anti-TEX12 | 1:50 | Ricardo Benavente (Biozentrum, University of Würzburg)<br>(Hamer et al., 2006) |
| Rabbit anti-SYCE3 | 1:2000 | Ricardo Benavente (Biozentrum, University of Würzburg)<br>(Schramm et al., 2011) |
| Rabbit anti-RAD51 | 1:500 | Millipore #PC130 |

Supplementary Table ST1. Antibodies used in this study.

### Supplementary Figure Legends

#### Supplementary Figure S1. SYCE2 microfoci and SYCP1 foci in *Syce3* spermatocyte nuclei.

(**A**) Distances between axial SYCE2 microfoci and their nearest SYCP1 foci. The distance between the centroid of each SYCE2 microfocus that overlaps the SYCP3 axis and the centroid of its nearest SYCP1 focus are shown alongside shuffled datasets assigning SYCE2 centroids to random positions on the SYCP3 axis for twenty repetitions. Crossbars represent quartiles; *, p<0.01, (Mann-Whitney U test, paired test used to compare observed with shuffled datasets, medians are 266.3, 316.8, 566.7 and 881.3 nm, n=25, 24 nuclei); 3 animals analysed for each *Syce3* genotype. Summary statistics are in Supplementary Table ST2.

(**B**) Scatter plot of axial SYCP1 foci areas in $Syce3^{\Delta/\Delta}$ and $Syce3^{WY/W}$ asynapsed pachytene nuclei. The 150 nm threshold distinguishing large SYCP1 foci (pink) is shown as a dotted line and the percentage of SYCP1 foci above this threshold is indicated for each genotype. *, p < 0.01 (Fisher's exact test, n=5567, 1402 foci). Summary statistics are in Supplementary Table ST2.

#### Supplementary Figure S2. RAD51 foci in *Syce3* and *Sycp1* mutant spermatocytes.

(**A**) Axial RAD51 foci frequency in asynapsed pachytene *Syce3*$^{\Delta/\Delta}$, *Syce3*$^{WY/WY}$ and *Sycp1*$^{-/-}$ nuclei. Crossbars represent quartiles; *, $p< 0.01$; ns, not significant (Mann-Whitney U test, medians are 196, 236 and 243 foci, n=39, 36, 34 nuclei); 3 *Syce3*$^{\Delta/\Delta}$, 3 *Syce3*$^{WY/WY}$ and 2 *Sycp1*$^{-/-}$ animals analysed. Summary statistics are in Supplementary Table ST2.

(**B**) SIM super-resolution images of *Sycp1*$^{+/-}$ pachytene spermatocyte chromosome spread immunostained for SYCP3 (blue), RAD51 (magenta) and SYCE2 (yellow). The region indicated with the dotted square is shown in the right-hand panels, RAD51 flares emanating from the axes are indicated by asterisks. Scale bar, 10 µm.

#### Supplementary Figure S3. Distances from SYCP3 axis traces in control synapsed pachytene and *Syce3* and *Sycp1* mutant asynapsed pachytene spermatocytes.

(**A**) Distance between paired SYCP3 axis traces in fully synapsed C57BL/6J wild type pachytene chromosome spreads. The distance from each point on the traced SYCP3 axis to its nearest point on its partner axis trace was measured. Gray dotted lines indicate 130 nm and 300 nm, green dotted line indicates 200 nm. Data represent 113515 data points from 14 nuclei; 3 animals analysed. Summary statistics are in Supplementary Table ST2.

(**B**) Violin plot showing the distribution of mean distances between each point on the skeletonized SYCE2 axis and their nearest points on traced, paired SYCP3 axes in fully synapsed pachytene C57BL/6J spermatocyte nuclei. Crossbars represent quartiles, green dotted lines represent 140 and 80 nm, gray dotted line represents 100 nm. Data represent 28323 data points from 11 nuclei; 3 animals analysed. Summary statistics are in Supplementary Table ST2.

(**C**) Violin plot showing the distribution of distances between axis-associated SYCE2 microfoci centroids and their nearest points on traced, paired SYCP3 axes in *Syce3* and *Sycp1* mutant asynapsed pachytene nuclei. Gray dotted line represents 200 nm. Data represent 5127, 3102 and 4974 data points from 24, 24 and 24 nuclei; 3 *Syce3*$^{\Delta/\Delta}$, 3 *Syce3*$^{WY/WY}$ and 2 *Sycp1*$^{-/-}$ animals analysed. Each of these mutants has an enriched population of SYCE2 microfoci located ~80-100 nm from the SYCP3 axis. Summary statistics are in Supplementary Table ST2.

#### Supplementary Figure S4. Uncropped gels for biochemical interactions between SYCP1 and central element components.

(**A-E**) Uncropped gels for amylose pulldowns shown in Figure 6A in which MBP-SYCP1N (1-362) and free MBP were mixed with (**A**) SYCE3, (**B**) TEX12, and the structural cores of (**C**) SYCE1, (**D**) SYCE1-SIX6OS1 and (**E**) SYCE2-TEX12.

(**F**) Uncropped gel of Figure 6B, corresponding to amylose pull-down of SYCE2-TEX12 following recombinant co-expression with MBP-SYCP1 constructs (as indicated) and free MBP (empty).

#### Supplementary Figure S5. SYCE2 is recruited to leptotene chromosomes independently of SYCP1.

(**A**) SIM super-resolution images of *Sycp1*$^{+/-}$ and *Sycp1*$^{-/-}$ leptotene spermatocyte chromosome spreads immunostained for SYCP3 (blue), RAD51 (magenta) and SYCE2 (yellow). Co-localised SYCE2 and RAD51 foci are indicated with asterisks. The regions indicated with the dotted square are shown at higher magnification in the right-hand panels. Scale bar, 10 µm.

(**B**) Distance from the centroid of each SYCE2 microfocus to the nearest SYCP3 axis mask in *Sycp1*$^{+/-}$ and *Sycp1*$^{-/-}$ leptotene spermatocyte nuclei. Shuffled data were obtained by assigning all SYCE2 microfoci in each nucleus to random locations within the nucleus over twenty iterations. Blue and red dotted lines represent 140 and 35 nm. Crossbars represent quartiles; *, $p< 0.01$ (Mann-Whitney U test, paired test used to compare observed with shuffled datasets, medians are 316, 703.6, 133.8 and 775.5 nm, n=10 nuclei); 2 animals analysed for each *Sycp1* genotype. Summary statistics are in Supplementary Table ST2.

#### Supplementary Figure S6. Overlapping images used to capture chromosome spreads.

(**A**) *Syce3*$^{PAM/PAM}$ images used to capture the chromosome spreads shown in Figure 2A. Overlapping images were taken with 15% overlap and stitched using Nikon NIS-Elements. Scale bar 10 µm. This chromosome spread was also stained with anti-SYCP1 antibodies (shown in Crichton et al., 2022).

(**B**) *Sycp1*$^{+/-}$ and *Sycp1*$^{-/-}$ images used to capture the chromosome spreads shown in Figure 3E. Overlapping images were taken with 15% overlap and stitched using Nikon NIS-Elements. Scale bar 10 µm.

(**C**) *Sycp1*$^{+/-}$ and *Sycp1*$^{-/-}$ images used to capture the chromosome spreads shown in Figure 4C. Overlapping images were taken with 15% overlap and stitched using Nikon NIS-Elements. Scale bar 10 µm.

(**D**) *Sycp1*$^{+/-}$ and *Sycp1*$^{-/-}$ images used to capture the chromosome spreads shown in Figure S6A. Scale bar 10 µm. Overlapping images were taken with 15% overlap and stitched using pairwise stitching in Fiji.

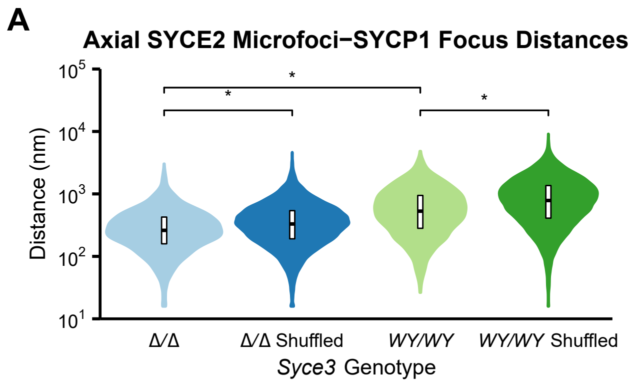
A
Axial SYCE2 Microfoci−SYCP1 Focus Distances
Distance (nm)
10^5
10^4
10^3
10^2
10^1
*
*
*
Δ/Δ
Δ/Δ Shuffled
WY/WY
WY/WY Shuffled
Syce3 Genotype

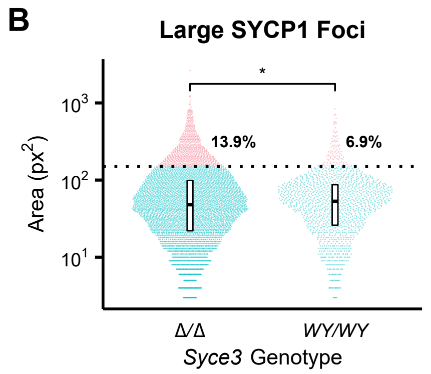
B
Large SYCP1 Foci
Area (px^2)
10^3
10^2
10^1
*
13.9%
6.9%
Δ/Δ
WY/WY
Syce3 Genotype

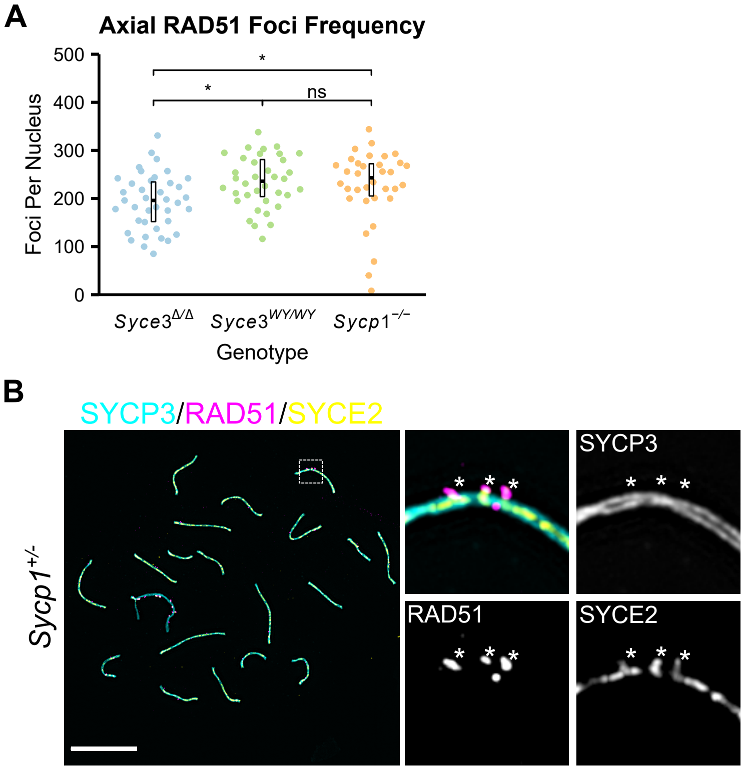
A
Axial RAD51 Foci Frequency
500
400
300
200
100
0
Foci Per Nucleus
*
*
ns
Syce3Δ/Δ
Syce3WY/WY
Sycp1−/−
Genotype
B
SYCP3/RAD51/SYCE2
Sycp1+/-
SYCP3
RAD51
SYCE2

Supplementary Figure S3

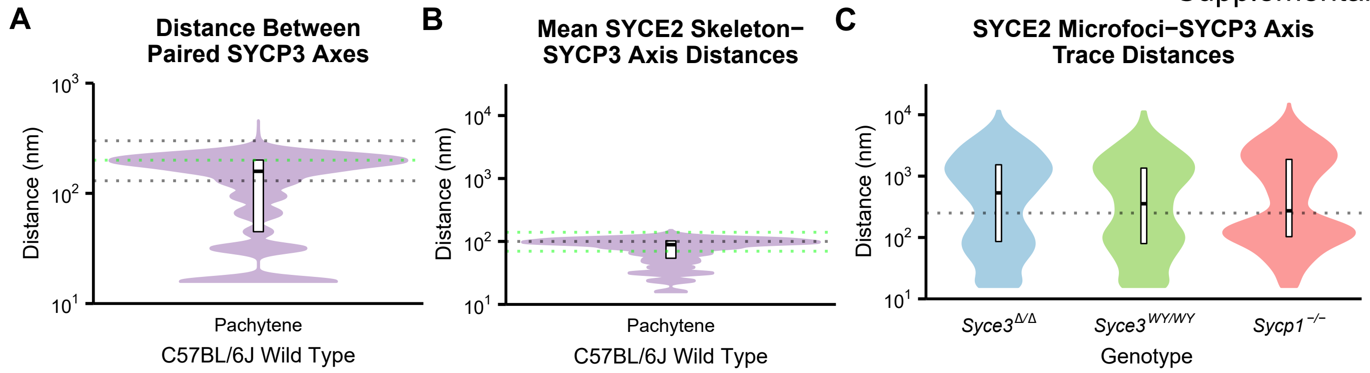

Supplementary Figure S4

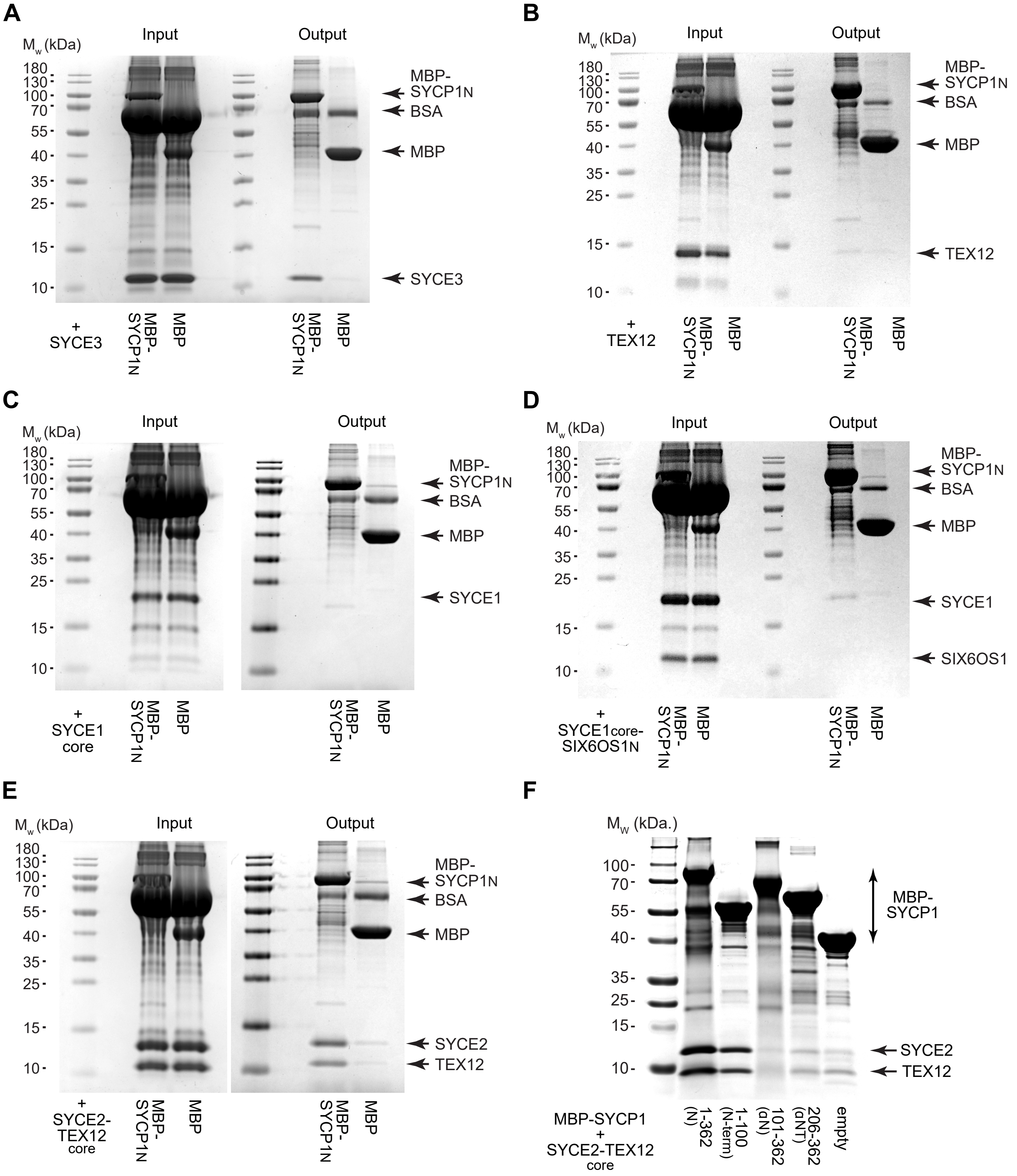

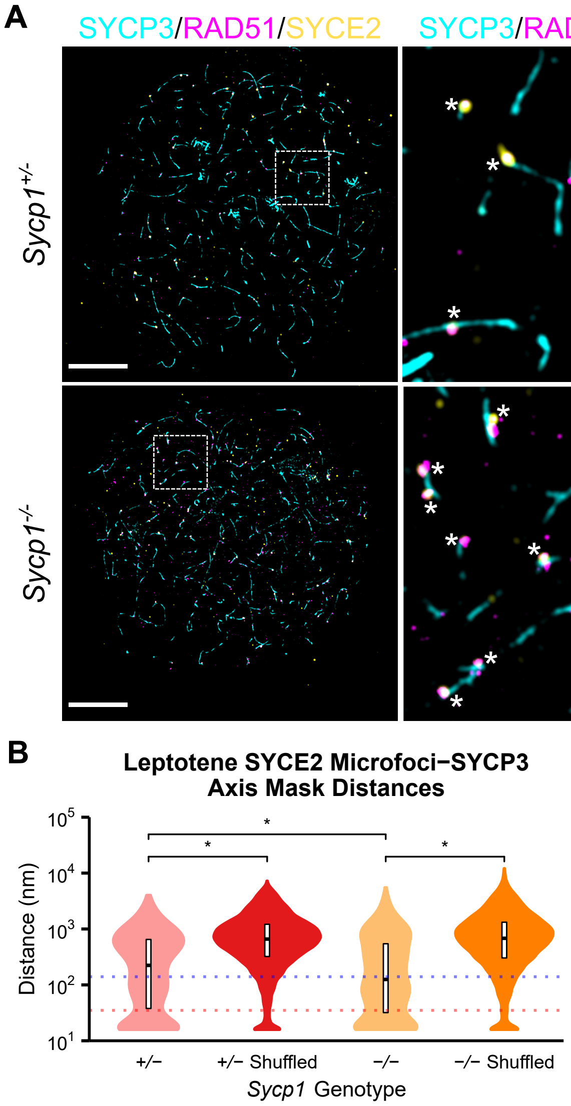
A
SYCP3/RAD51/SYCE2
SYCP3/RAD51/SYCE2
SYCP3/RAD51
SYCP3/SYCE2
Sycp1+/-
Sycp1-/-
B
Leptotene SYCE2 Microfoci–SYCP3
Axis Mask Distances
Distance (nm)
10^5
10^4
10^3
10^2
10^1
+/–
+/– Shuffled
–/–
–/– Shuffled
Sycp1 Genotype

### Supplementary Figure S6

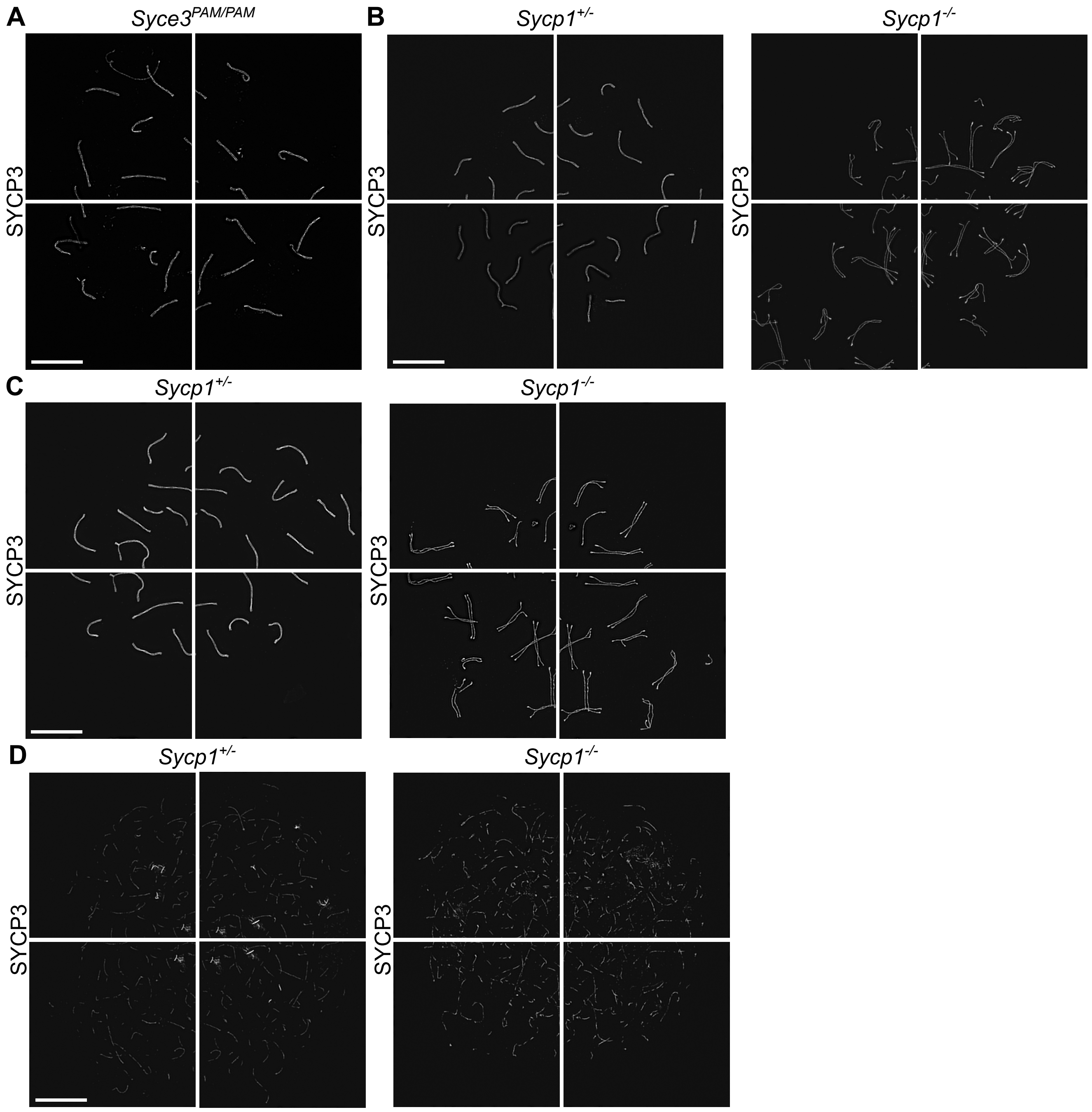
